# Transcriptional control of the *lacZ* promoter is under directional and diversifying selection

**DOI:** 10.1101/2022.03.17.484780

**Authors:** Markéta Vlková, Olin K. Silander

**Affiliations:** School of Natural and Sciences, Massey University, Auckland, New Zealand

## Abstract

Bacterial cells often respond to changes in the environment by modifying protein expression. This can be achieved through changes in transcriptional or translational activity, or both. Recent research has shed some light on how natural selection shapes overall protein expression. Still, little is known about how selection acts on transcription or translation individually. To address part of this question, we implement an experimental system which allows us to measure how genetic changes affect transcription only, excluding the effects on translation. We use this system to quantify changes in three regulatory phenotypes of the *lacZ* promoter: transcriptional activity, plasticity, and cell-to-cell variability. We compare these phenotypes from segregating variants that have been subject to natural selection, and random variants that have never been subjected to natural selection. We show that natural selection filters out mutations causing large changes in transcriptional levels from the *lacZ* promoter. Further, we detect directional selection acting on transcriptional plasticity in combinations of glucose, galactose and lactose environments. Focusing on cell-to-cell variability in transcription, we describe both directional and diversifying selection acting on this phenotype depending on the environment used. We also observe a link between the phylogeny of the environmental *E. coli* strains and high and low transcriptional noise levels in glucose which are mediated by just one or two SNPs. Our results thus provide new insight into how one of the most well-characterized bacterial promoters is shaped in nature by selection.

## Introduction

Bacteria respond to dynamically changing environmental conditions by altering their physiology and metabolism. Natural selection acts on aspects of these responses, such as the level, speed, sensitivity, and variability (Silander et al. 2012; Keren et al. 2016; Hodgins-Davis et al. 2019; Schmiedel et al. 2019; Hawkins et al. 2020; Vlková and Silander 2021). Frequently these aspects are controlled by mediating protein levels, which in turn are mediated by changes in transcriptional and translational activity.

However, there is little data on whether over evolutionary time, genetic variation in transcriptional or translational processes are most responsible for generating new phenotypic variation and mediating phenotypic change, i.e. changes in protein levels and the downstream cellular responses. It is possible that most new mutations affect phenotypes by altering transcription but not translation. If this were the case, under stabilizing selection we would expect that most new mutations that have large phenotypic effects would be filtered by selection due to their effect on transcription. However, the converse could also be true: most new mutations could affect phenotypes by altering translation.

The phenotypic effects of new mutations, and the relative dominance of transcriptional or translational effects may also depend on the specific regulatory context. For example, from an energetic perspective, it seems straightforward that in order to produce a specific level of protein, it is optimal to produce as little mRNA as possible while still producing sufficient protein - minimizing transcription level and maximizing translation (Frumkin et al. 2017). While this is still an example of stabilizing selection on the phenotype (expression level), at such regulatory extremes, the phenotypic effects of new mutations might be expected to often increase transcription and decrease translation. Because of this bias in mutational effects, selection would appear far more directional when considering each mechanism alone (e.g. selection to minimize transcription levels).

However, there are well-established trade-offs in using a low transcription-high translation solution: low transcription rates can slow down the response time to an external environmental signal, and increase variability between isogenic cells in their response (Maeda and Sano 2006; Sayut et al. 2007; Taniguchi et al. 2010). Again, the result of such a scenario might be selection for intermediate levels of transcription and translation, with apparent stabilizing selection on both.

Overall, it is unclear what mechanisms are most responsible for generating new variation in expression phenotypes, and how regulatory changes are generally mediated over evolutionary time (mostly via transcription, translation, or a combination of both). Nor is it readily apparent which evolutionary forces are acting on regulatory phenotypes (e.g. directional, stabilizing, or diversifying). While the mechanisms by which new variation in expression phenotypes is generated is important from a mechanistic evolutionary perspective, there are also implications for more applied contexts. An understanding of whether new phenotypes are more easily generated by changing transcription or translation would have applications in synthetic biology. Although the evolutionary forces shaping regulatory phenotypes in natural populations are not fully understood, it is readily apparent that changes in regulation can evolve rapidly (Gresham et al. 2008; Tenaillon et al. 2016; Yona et al. 2018; Khademi et al. 2019) via very few mutations (Metzger et al. 2015; Duveau et al. 2017; Vlková and Silander 2021). There are a number of mechanisms by which this can occur. The frequency of transcription can be changed by increasing or decreasing the strength of transcription factor binding sites, either activators or repressors. Alternatively, the strength of the sigma factor binding site can be adjusted through changes in the nucleotide sequence of the -10 and -35 elements, or the distance between them (Kennell and Riezman 1977; Iyer and Struhl 1996; Browning and Busby 2016). Changes to translational regulation can also affect downstream protein levels, and these can occur through adjustments to the binding energy of the ribosomal binding site, including specific binding of antisense RNAs, the stability and folding energy of the 5’ end of the mRNA, or by changing codon content, which may affect ribosomal speed or ribosomal pausing (Kudla et al. 2009; Naville and Gautheret 2009; Desnoyers et al. 2013; Whitaker et al. 2015; Mustaev et al. 2017; Bailey et al. 2021).

Here we focus on the evolution of regulatory responses and the associated mechanisms for a canonical example of physiological adaptation, the *E. coli lac* operon. This operon is involved in the metabolism of lactose, and is regulated by the presence of lactose in the environment and cAMP in the cell (Hudson and Fried 1990). When glucose levels are high, the levels of free cAMP drop, inhibiting transcription from the *lac* operon; when glucose levels are low and lactose levels are high, then the operon is activated (Jobe and Bourgeois 1972; Wanner et al. 1978; Kuo et al. 2003; Wheatley et al. 2013). The expression is largely dependent on the levels of intracellular cAMP and lactose (Ozbudak et al. 2004; Kuhlman et al. 2007).

However, most of what is known about the *lacZ* promoter activity has been obtained using classical laboratory strains. It has been shown that there is a substantial genetic and phenotypic variability in *lacZ* promoter among environmental isolates of *E. coli* (Phillips et al. 2019; Vlková and Silander 2021). To understand the regulatory effects of new mutations and segregating polymorphisms, here, we implement an experimental system that allows measurement of transcriptional effects alone. This enables us to test whether many of the phenotypic differences between isolates are dependent solely on regulatory changes in translation, or whether they involve changes to transcriptional regulation.

An unexpected benefit of the experimental system we use here is that it has substantially increased sensitivity for detecting changes in transcriptional regulation. Thus, we can detect a wider range of evolutionary forces acting on transcriptional regulation, including both directional and diversifying selection. Using this system, we observe directional selection for low transcriptional noise in lactose and high plasticity in combinations of glucose and lactose.

Furthermore, we detect a long-term segregating polymorphism that appears to affect the level of transcriptional noise from *lacZ* promoter variants in non-lactose environments. This work thus yields new insight into how the activity of one of the most well-characterized bacterial promoters is shaped in nature.

## Results

### 1. A plasmid based approach to infer the action of selection on transcriptional control

In order to investigate the selection pressure acting on the transcriptional regulation of the *lacZ* promoter in *E. coli*, we first quantified genetic diversity in the regulatory region of the *lac* operon in a collection of 135 environmental *E. coli* isolates (Ishii et al. 2006). Here, we have designated the *lacZ* promoter as the entire *lacI-lacZ* intergenic region (IGR) as well as 88 bp of the 3’ end of the upstream *lacI* ORF and 69 bp of the 5’ end of the downstream *lacZ* ORF (**Fig. 1a**). We include these regions, as ORFs that flank IGRs are known to frequently affect regulation of gene expression (Zaslaver et al. 2006; Vlková et al. 2021). We identified 26 *lacZ* promoter variants segregating in this set of environmental *E. coli* isolates (**Supplementary Figure 1** and **Supplementary Table 1**). These polymorphisms were present both inside and outside transcription factor binding sites, and we observe no clear evidence for polymorphism clustering in segregating variants (**Supplementary Figure 1**).

**Figure 1:**
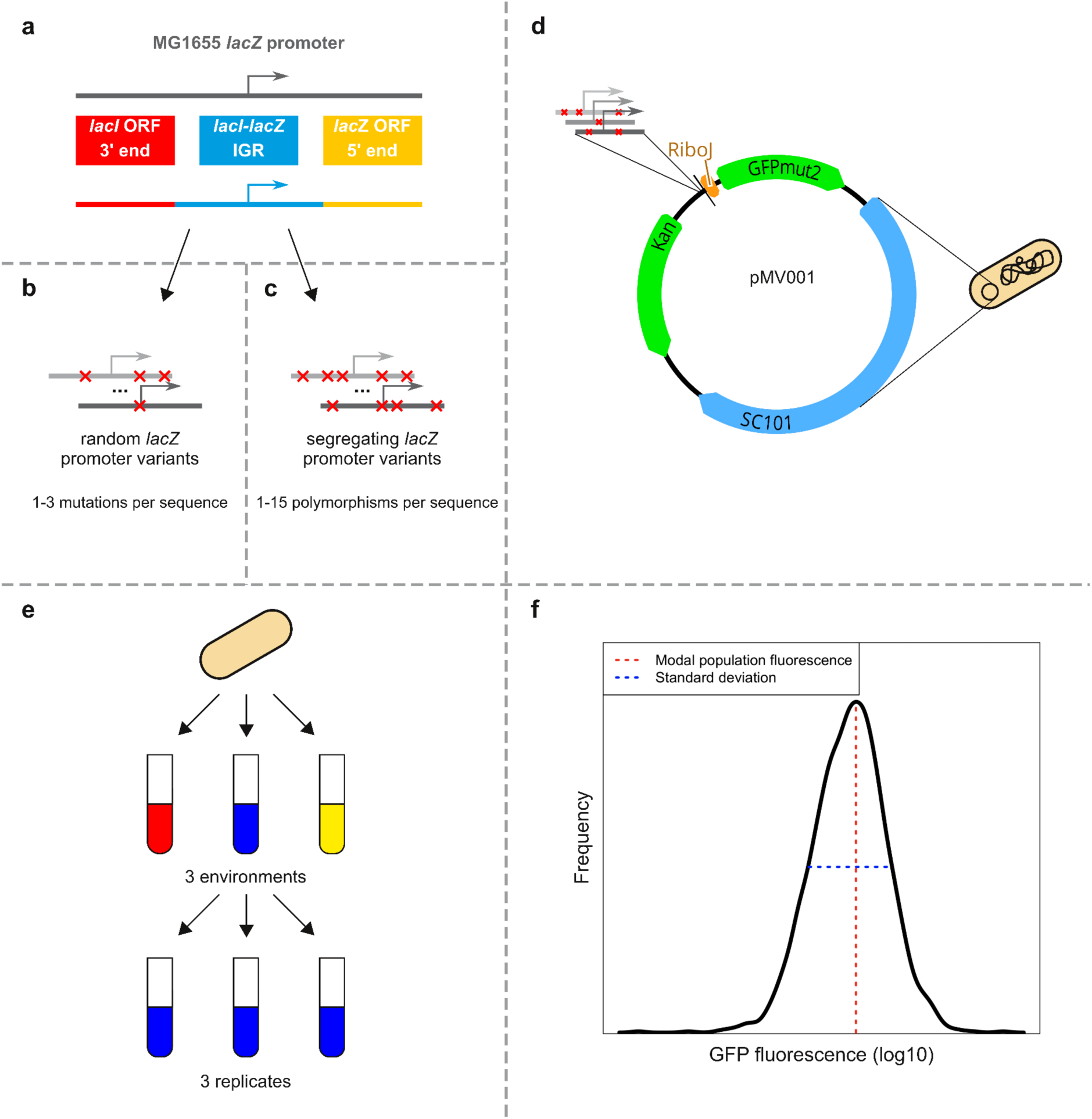
Experimental design to assay the effects of segregating polymorphisms and random mutations on transcriptional control from *lacZ* promoter. **a**, The sequences we refer to as *lacZ* promoter variants consist of the *lacI-lacZ* intergenic region (IGR), 88 bp of the 3’ end of the upstream *lacI* open reading frame (ORF), and 69 bp of the 5’ end of the downstream lacZ ORF. **b**, We performed random PCR mutagenesis using the MG1655 *lacZ* promoter variant as a template with a target mutation rate of 1.5 mutations per promoter variant. **c**, We also PCR-amplified variants of *lacZ* promoter segregating among environmental *E. coli* isolates from a DNA pool. The average number of polymorphisms across obtained segregating variants (as compared to MG1655) is almost 10-times higher than the average number of mutations for random variants. **d**, We cloned the resulting PCR amplicons (both segregating and random) upstream of GFPmut2 (Vlková et al. 2021). We Sanger sequenced all the promoter variants to confirm the presence and location of mutations. From mutagenesis, only the variants containing 1 to 3 mutations were used for further phenotypic assays. **e**, Each bacterial clone was cultured in media containing glucose, galactose, or lactose as the primary carbon source in triplicate. **f**, Using flow cytometry we quantified the modal population fluorescence and the standard deviation of population fluorescence.

This plasmid-based system omits the O2 operator of *lacZ* promoter which lies over 350 bp into the *lacZ* gene coding sequence. However, the effect of the O2 operator on expression from the *lacZ* promoter, this effect is very weak (Oehler et al. 1990). In addition, there is additional selection as it is within a coding region (Rech et al. 2014). For all these reasons, we have excluded it.

In nature, all newly generated variants (i.e. those arising through mutation or recombination) may be filtered by natural selection based on their phenotypic effects. Our goal here is to understand the strength of this filtering and the effects of selection on transcriptional regulation. However, many of the polymorphisms present in segregating promoter variants are downstream of the *lacZ* transcription start site, and thus present in the transcribed mRNA. These changes in the *lacZ* mRNA sequence may result in phenotypic changes by altering translation, for example by changing mRNA folding or ribosome initiation. In order to investigate selection forces acting on transcription in isolation from translation, we sought a method to disentangle phenotypic differences that are mediated by changes in transcription from phenotypic differences mediated by changes in translation.

To do this, we designed a system in which a GFP open reading frame is placed downstream of a promoter variant on a low-copy number plasmid (Zaslaver et al. 2006). We placed a self-cleaving ribozyme, RiboJ, upstream of the GFP open reading frame (Lou et al. 2012; Vlková et al. 2021) (**Fig. 1d**). When this mRNA is transcribed, RiboJ immediately cleaves and removes the 5’ end of the mRNA, including any polymorphisms or mutations that are present (**Fig. 2**). This cleavage is extremely rapid, and at any particular time, more than 95% of all mRNAs in a cell are cleaved. Thus, any observed effects of polymorphisms on GFP expression and fluorescence are necessarily mediated solely by changes in transcription (Vlková et al. 2021). This contrasts with mRNA molecules lacking RiboJ, as sequence changes present in these mRNAs may also affect GFP expression and fluorescence by affecting translation.

**Figure 2:**
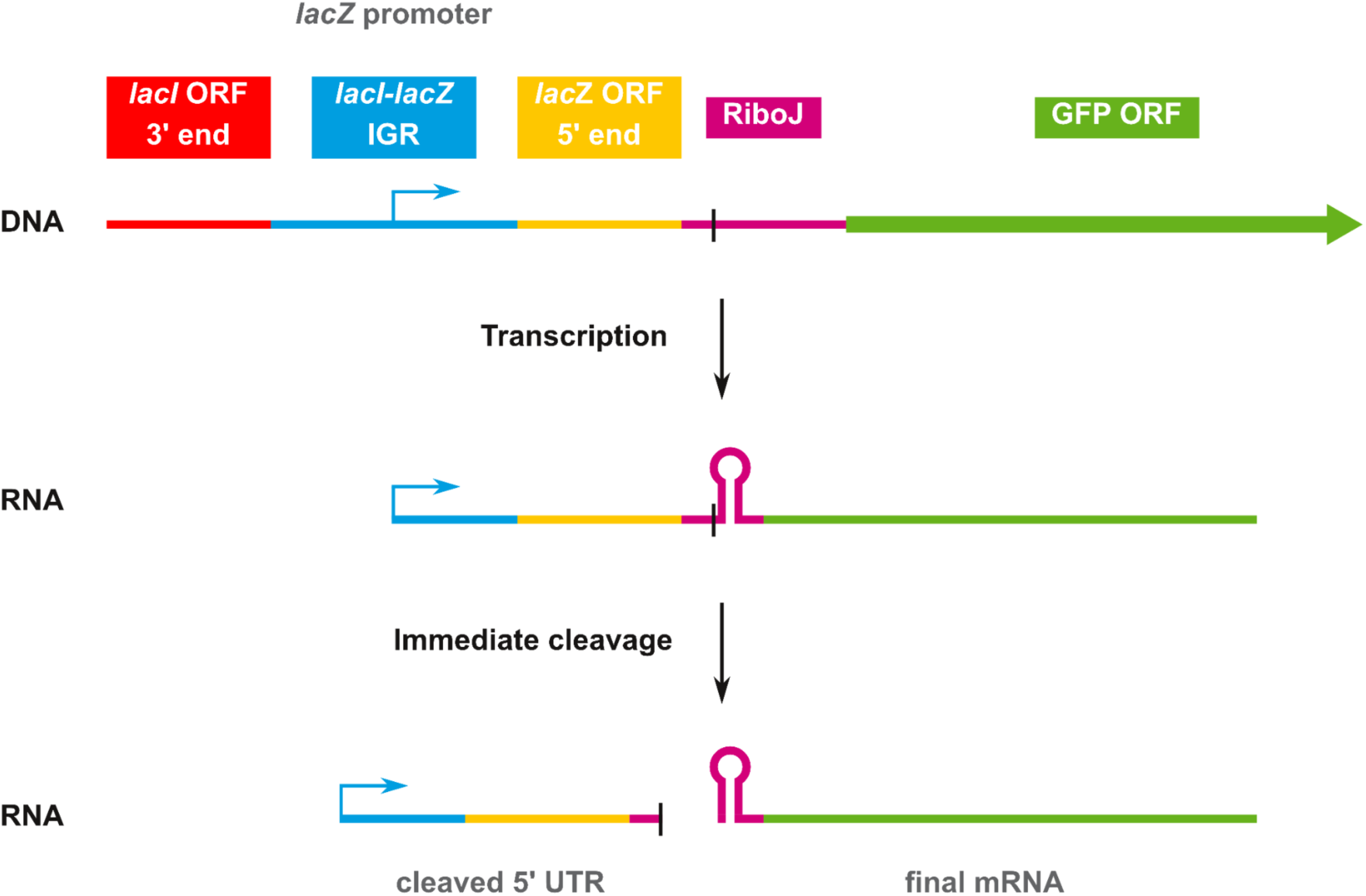
Removal of 5’ sequence variation via RiboJ ribozyme autocatalytic cleavage activity. The RiboJ sequence is placed between the *lacZ* promoter and GFP open reading frame (ORF). The arrow in the middle of the *lacI-lacZ* intergenic region (IGR) represents the transcription start site. When transcription is induced from the *lacZ* promoter, the RiboJ RNA sequence produces a strong secondary RNA structure resulting in fast and efficient cleavage at a specific site, cutting off most of the 5’ untranslated region (UTR). The final mRNAs that undergo translation from different *lacZ* promoter variants are thus identical, even when there are polymorphisms downstream of the transcription start site. The differences in GFP fluorescence measured from the promoter variants are the result of differences solely in transcription.

To understand the filtering process of natural selection on transcriptional regulation, we used PCR mutagenesis to generate a set of promoter variants with random mutations that have not been filtered by selection (**Materials and Methods**). By comparing the regulatory phenotypes of segregating and random promoter variants, we can infer how natural selection has acted. Note that we designate genetic differences in segregating promoter variants as “polymorphisms,” and genetic differences in the random promoter variants as “mutations” to differentiate genetic differences that have or have not been filtered by selection.

To generate this set of promoter variants unfiltered by selection, we took the MG1655 variant of the *lacZ* promoter (**Fig. 1a**) and performed PCR mutagenesis, generating approximately 1.5 mutations per promoter variant (**Fig. 1b** and **Supplementary Figure 2**). We cloned these promoters upstream of the GFP open reading frame on the plasmid-based system with RiboJ described above. At the same time, we cloned the 18 of the 26 segregating promoter variants into the same plasmid (**Fig. 1c**, **Fig. 1d**, **Supplementary Table 1**, and **Supplementary Table 2**). We sequenced all the random promoter variants, and selected 92 that contained between one and three mutations. We note that this random promoter library is considerably less diverse than the segregating variants: the average pairwise identity for all segregating variants is 96.25%, compared to 98.97% for the random variants.

We next quantified the regulatory phenotypes of each of the promoter variants in each of the libraries. We transformed all the promoter constructs into *E. coli* K12 MG1655 (**Supplementary Table 1** and **Supplementary Table 2**), and grew each clone in the presence of three different carbon sources (0.4% glucose, 0.4% galactose, and 0.4% lactose) for four hours, at which time the cultures were in the exponential growth (**Fig. 1e**). We then measured the amount of fluorescence from each cell using flow cytometry. We used the changes in fluorescence as a proxy for changes in transcriptional activity (Vlková et al. 2021). For each sample population we calculated two metrics of regulatory phenotypes, the modal population fluorescence and the standard deviation (**Fig. 1f**). All measures were taken from three full biological replicates (**Materials and Methods**).

We first compared the fluorescence levels we observed from promoter variants with RiboJ to the fluorescence levels of identical promoter variants lacking RiboJ that we had previously generated (Vlková and Silander 2021). We found that the fluorescence levels from the promoter variants in these two systems were highly correlated, with correlations of 0.86, 0.87, and 0.91 in the glucose, galactose, and lactose environments, respectively (Pearson’s rho; **Supplementary Figure 3**). We also observed an approximately ten-fold systematic increase in fluorescence levels when RiboJ was present. This increase was expected, and has been ascribed as resulting from the specific mRNA folding after the RiboJ self-cleaving (Carrier and Keasling 1997; Clifton et al. 2018; Neves et al. 2020). Thus, the addition of RiboJ brings two advantages. First, it allows us to determine the extent to which selection has acted on regulatory phenotypes mediated solely via transcriptional mechanisms. Second, the systematic increase in fluorescence in the presence of RiboJ increases the sensitivity of our measurements.

### 2. Directional selection acts on *lacZ* promoter activity in glucose and lactose

To infer the filtering effects of natural selection we compared the fluorescence levels of the random promoter variants to the segregating promoter variants, including the progenitor MG1655 variant. We expected that if there were systematic differences in fluorescence between the random and segregating sets of variants, that this is caused by selection tending to remove new variants that cause undesired changes in transcription. For example, if most random variants exhibit systematically higher fluorescence than segregating variants, this suggests that selection has removed newly generated variants with higher fluorescence levels.

We found that the segregating variants differed by approximately three-fold in transcriptional activity in each of the environments (**Fig. 3a**). While the majority of random variants also ranged approximately three-fold in transcriptional activity, a small number had more than ten-fold higher or lower transcriptional activity than the MG1655 variant from which they were derived.

**Figure 3:**
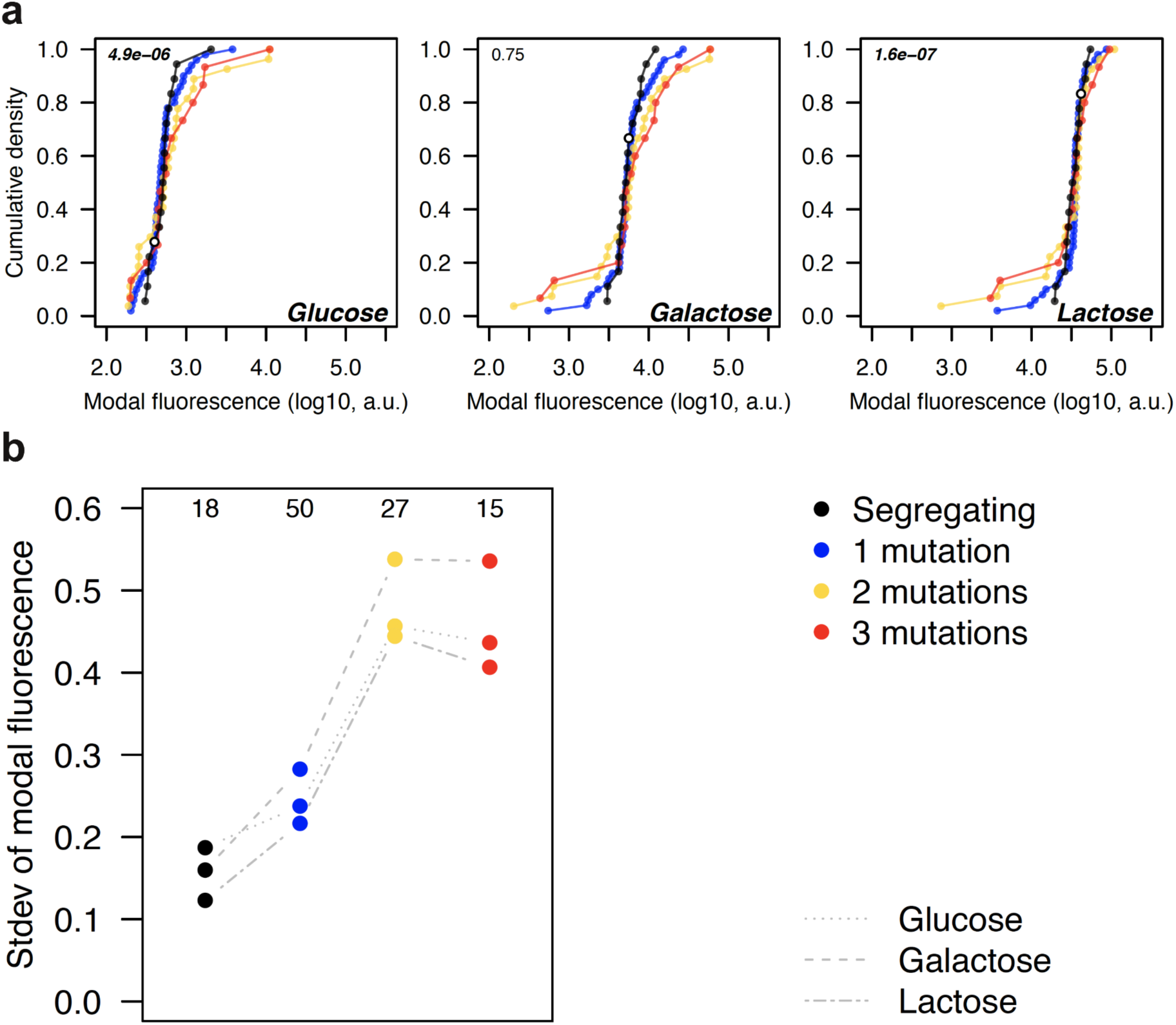
Selection acts to maintain similar transcription levels in segregating variants. **a**, Cumulative density of modal population transcriptional levels of random and segregating *lacZ* promoter variants in the MG1655 genetic background in all environments (glucose, galactose, and lactose). Random variants are stratified by the number of mutations they differ from the MG1655 variant from which they were derived. Segregating variants include the MG1655 variant which is displayed as a white circle in each plot. The numbers in the top-left corner represent p-values from the two-sided binomial test indicating whether random variants are as likely to increase as decrease transcriptional level when the MG1655 variant is randomly mutated. **b**, Standard deviation (stdev) of modal population fluorescence values for each promoter variant group in all environments. The standard deviation generally rises with an increasing number of mutations among random variants. The segregating variants have among the lowest standard deviations while having up to 15 polymorphisms relative to the MG1655 variant. The numbers above each group of points indicate the sample sizes.

However, segregating variants are approximately four-fold more genetically diverse. We stratified the random variants according to the number of random mutations they had compared to the progenitor MG1655 promoter variant. We found that as the number of random mutations in a promoter increased, their transcriptional activity diverged more and more from the transcriptional activity of the segregating variants (**Fig. 3b**).

Furthermore, the divergence in transcriptional activity was not symmetric across environments. Specifically, we found that in the glucose environment, in which the *lac* operon is highly repressed, the vast majority of random variants increased transcriptional activity. We calculated the probability of the observed distribution of effects to a distribution symmetric around the activity of the MG1655 variant, and found that the fraction of random variants that increased activity was much greater than expected (68 out of 92, p = 4.9e-06, two-sided binomial test). In contrast, in the lactose environment, in which the *lac* operon is highly active, the majority of random variants decreased their transcription (71 out of 92, p = 1.6e-07, two-sided binomial test). We found no difference in the galactose environment (p = 0.75). However, we note that these results are highly dependent on the precision of the fluorescence levels that we have measured for the MG1655 variant, although these are likely to be accurate as we performed all reported measurements in triplicate.

In addition, in all environments, we found that segregating variants exhibited lower phenotypic variation compared to random variants (**Fig. 3b**) although these results were not significant (p = 0.07 for glucose; p = 0.10 for galactose; p = 0.80 for lactose, Fligner-Killeen test of homogeneity of variances). This is despite segregating variants harboring much higher levels of genetic variation. These data thus suggest that selection has acted to remove mutations that have large effects on transcriptional activity. Moreover, the results in the glucose environment suggest that in MG1655, selection has acted to reduce transcriptional activity, as the majority of random variants increase activity. Conversely, in MG1655, selection has acted to maximize transcriptional activity in the lactose environment.

### 3. Selection on transcriptional plasticity

Regulating transcription such that it is maximized in one environment and minimized in a second can be viewed as maximizing transcriptional plasticity - the size of the change in transcriptional activity is dependent on environmental conditions. To visualize plasticity in two environments, we can plot the fluorescence level in one environment versus the fluorescence level in the second environment, with the x = y isocline indicating equal fluorescence and thus equal transcriptional activity in both environments. We quantified plasticity as the shortest distance from this isocline (**Fig. 4a**). Thus, promoter variants that lie directly on this line have equal transcriptional activity in both environments and no plasticity; the further they are from this line, the higher their plasticity. When considering three environments, we quantified plasticity using an analogous measure: the shortest distance to the isocline from a point defined by the fluorescence levels in all three environments considered here: glucose, galactose, and lactose (**Fig. 4c**). Note that plasticity calculated from the combination of all three environments (**Fig. 4d**) strongly mirrors the pattern of plasticity from the combination of glucose and lactose only (**Fig. 4b**, Glu:Lac). This is due a strong dependence of the plasticity from three environments on the largest difference in fluorescence levels between any two environments, which in the case of *lacZ* promoter variants are glucose and lactose, as expected.

**Figure 4:**
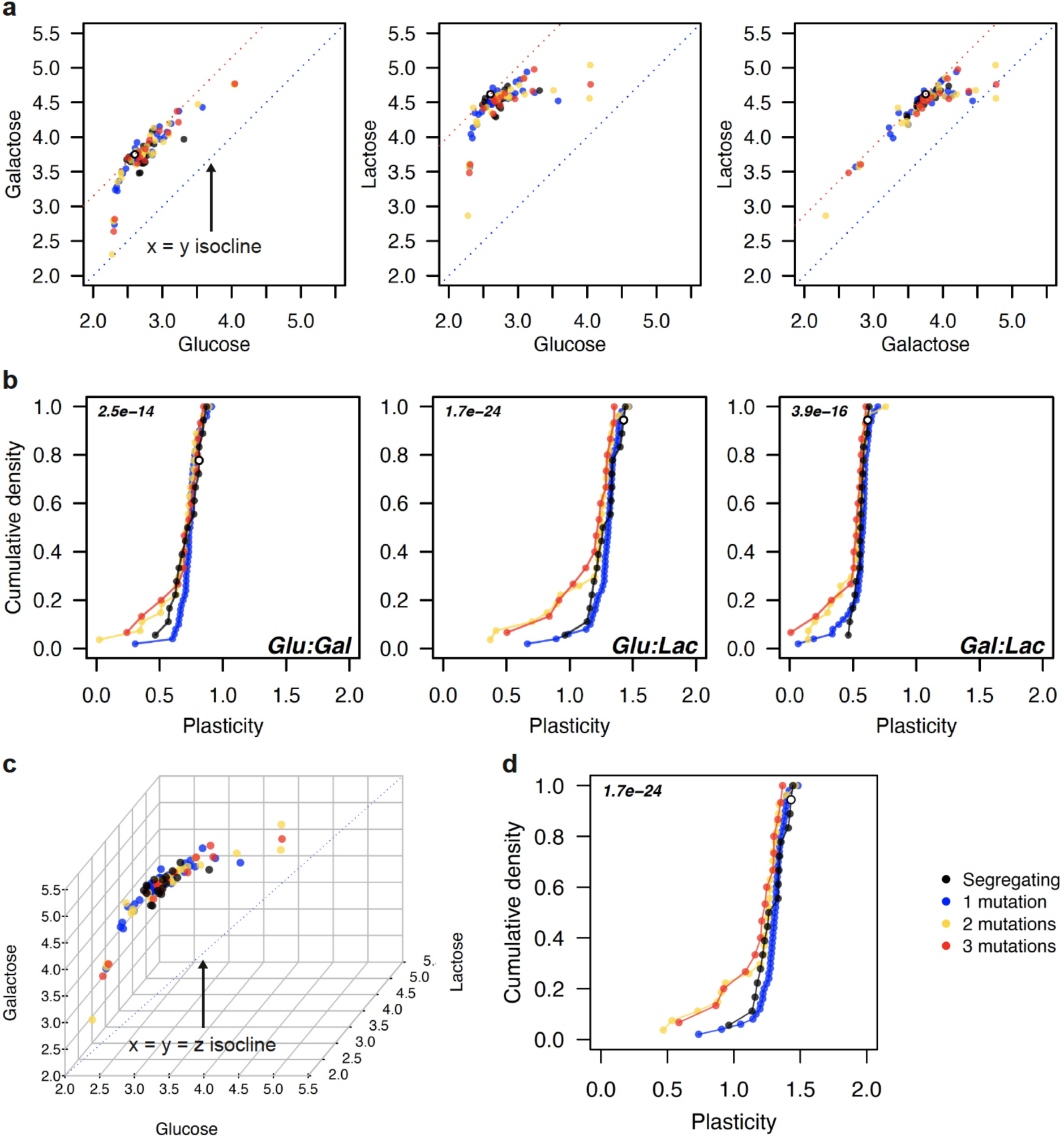
Directional selection acts to maintain high transcriptional plasticity in segregating variants. **a**, Comparison of modal fluorescence levels from random and segregating variants in pairs of environments as indicated by the x and y axis labels. The plasticity from a pair of environments is calculated as the distance of a datapoint (promoter variant) from an isocline of equal fluorescence levels (x = y, blue dotted line). Promoter variants lying on the red dotted line have plasticity levels equal to the MG1655 variant through which the line runs. **b**, Cumulative plasticity levels from random and segregating variants in pairs of environments. **c**, Comparison of modal fluorescence levels from random and segregating variants in all three environments. The plasticity from three environments is calculated as the distance of a datapoint (promoter variant) from an isocline of equal fluorescence levels (x = y = z, blue dotted line), analogous to the approach in **a**. **d**, Cumulative transcriptional plasticity levels from random and segregating variants in all three environments. Across all panels the random variants are stratified by the number of mutations they differ from the MG1655 variant from which they were derived. Segregating variants include the MG1655 variant which is displayed as a white circle. The numbers in the top-left corner in panels **b** and **d** represent p-values from the two-sided binomial test indicating whether random variants are as likely to increase as decrease plasticity when the MG1655 variant is randomly mutated.

We found that almost all random variants had reduced plasticity compared to MG1655 (81 out of 92, p = 2.5e-14 for glucose vs. galactose; 90 out of 92, p = 1.7e-24 for glucose versus lactose; 83 out of 92, p = 3.9e-16 for galactose vs. lactose, two-sided binomial test, **Fig. 4b**). Together with the results above, this suggests that for MG1655, selection has acted to increase plasticity by decreasing transcription in glucose and increasing it in lactose.

Relative to MG1655, we found that segregating variants generally exhibited lower plasticity, and in fact did not differ in plasticity compared to the random variants (p = 0.75 for glucose vs. galactose; p = 0.48 for glucose vs. lactose; p = 0.81 for galactose vs. lactose; p = 0.47 for all three environments, two-sided Wilcoxon rank-sum test). However, random promoters with more than two mutations exhibited a long tail of lower plasticity (**Fig 4b**).

Previously, we showed that relative to other promoters in *E. coli*, segregating variants of the *lacZ* promoter vary the most in expression levels within single environments (Vlková and Silander 2021). The absence of strong differences between segregating and random variants in plasticity support these results, indicating that selection on transcriptional activity and plasticity in the *lacZ* promoter is relatively relaxed, and new mutations are not strongly filtered by natural selection. An alternative explanation for this observation is that plasticity may be highly dependent on the genetic background; thus, the plasticity that we have measured in MG1655 for each of the segregating variants may be lower than would be observed in the native background of the variants.

### 4. Both diversifying and directional selection act on transcriptional noise

Finally, we tested whether selection has acted to filter mutations according to their effects on transcriptional noise. Here, “transcriptional noise” refers to the cell-to-cell variability in transcriptional activity within an isogenic population. As the variation in fluorescence is highly dependent on the modal fluorescence level, we quantified the level of transcriptional noise by calculating the vertical deviation from a spline fitted to modal fluorescence versus the standard deviation divided by the modal fluorescence, a measure analogous to the coefficient of variation and which we refer to as the modal CV, or mCV (Vlková and Silander 2021). This measure of noise is independent of fluorescence level. In addition, because the variation in population fluorescence is highly dependent on the growth environment, we fit a spline for all promoter variants in each environment. This allows us to infer how selection is acting on transcriptional noise within a specific growth environment, rather than between environments, for which other selective factors (e.g. selection on transcriptional activity) may overwhelm selection on noise.

With this noise metric, variants with a higher mCV than expected given their modal fluorescence value have “high” noise, while variants with lower mCV than expected have “low” noise. As noted above, this metric also results in a different value of noise being calculated for each promoter variant in each environment (**Fig. 5a**).

**Figure 5:**
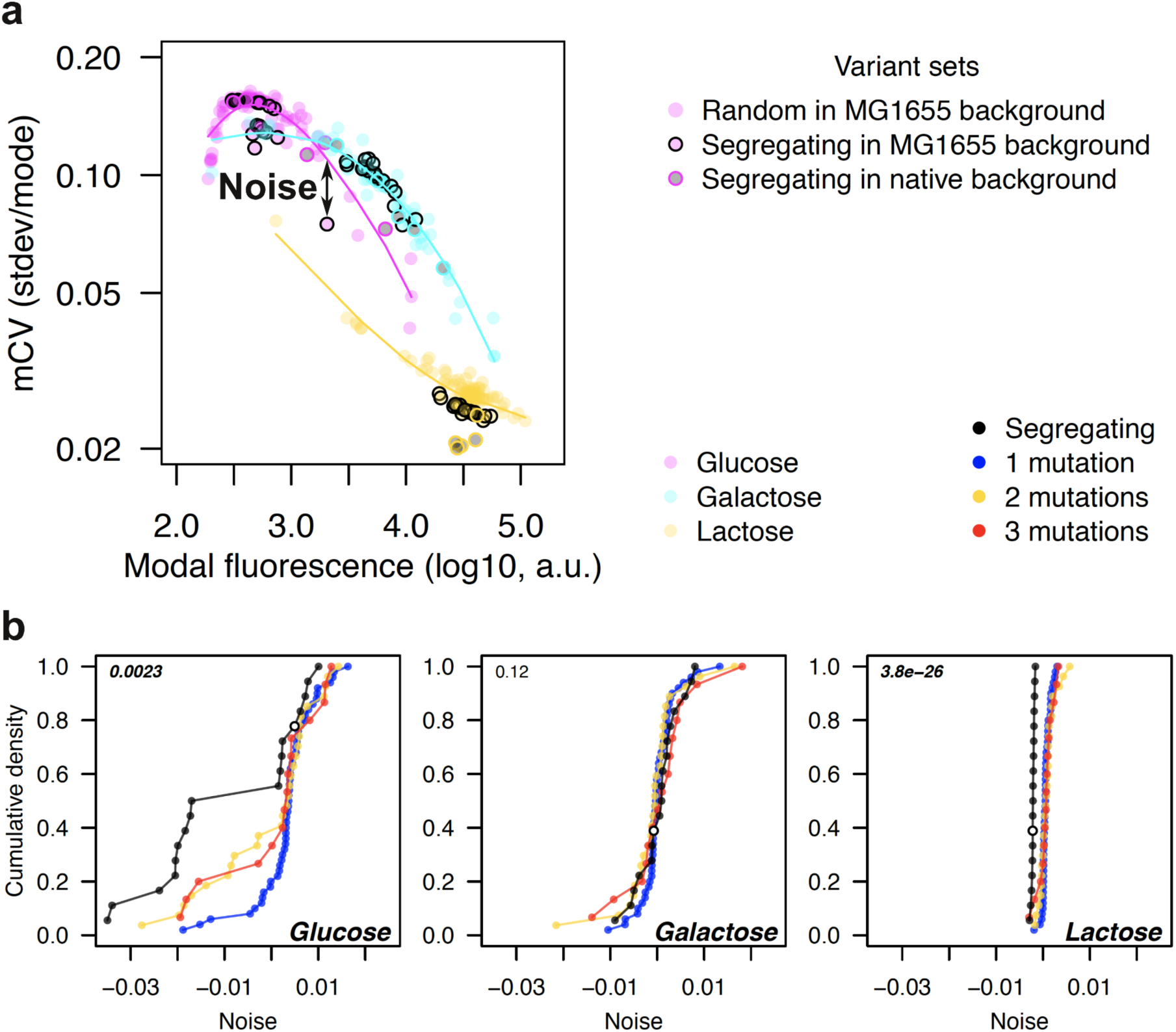
Selection on transcriptional noise levels in segregating *lacZ* promoter variants. **a**, We fit smoothing splines to modal population fluorescence vs. the standard deviation of population fluorescence normalized by modal fluorescence (mCV) for all variants (segregating and random) in all genetic backgrounds. We fit a separate spline for each environment. The term “noise” is used for the vertical deviation of each variant, i.e., deviation in the mCV from the fitted spline as indicated by the arrow. **b**, Cumulative distributions of transcriptional noise from random and segregating variants. Random variants are stratified by the number of mutations they differ from the MG1655 variant from which they were derived. Segregating variants include the MG1655 variant, which is displayed as a white circle. The numbers in the top-left corner represent p-values from a two-sided binomial test indicating whether random variants are as likely to increase as decrease transcriptional noise.

We first considered differences in transcriptional noise between segregating variants and random variants in lactose, where we expect full transcriptional activation. We found that all except for one of the 92 random variants (derived from the MG1655 promoter variant) exhibited higher levels of noise than the MG1655 variant (**Fig. 5b**). The MG1655 variant exhibited similar noise levels as the other segregating variants, and considering all segregating variants together, we found that they had significantly lower levels of transcriptional noise compared to random variants (median of -0.0021 ± 0.0003 for segregating variants vs 0.0005 ± 0.0013 for random variants, p = 9.4e-11, two-sided Wilcox rank-sum test). This suggests that in natural populations almost all mutations that detectably increase transcriptional noise when lactose is the primary carbon source are filtered out, indicative of selection for low transcriptional noise in lactose.

In strong contrast, in glucose, where we expect almost full repression of the *lac* operon, the majority of random variants had lower levels of noise than their MG1655 progenitor (61 out of 92, p = 2.3e-03, two-sided binomial test; **Fig. 5b**). In addition, we found that transcriptional noise varied considerably between segregating variants, with nine segregating variants, including MG1655, exhibiting high noise (between 0 and 0.01), and nine exhibiting low noise (all less than -0.015, **Fig 5b**). Surprisingly, a small number of the random variants had noise levels that approached those of the low-noise segregating variants. This suggests that low levels of noise are easy to achieve from a mutational standpoint, but are not always selected for. This might be due to most low noise phenotypes having deleterious effects on other regulatory phenotypes (transcriptional activity, plasticity, or both). It is also possible that for some isolates, high noise is advantageous when glucose is the primary carbon source.

Finally, in galactose, we found no strong differences in noise levels between the random and segregating variants, although the random variants followed a similar pattern to that observed for other phenotypes, as variants with greater genetic divergence from MG1655 also exhibited great phenotypic divergence. This is apparent in the tails of the distributions, which are dominated by the random mutants with two and three mutations relative to the progenitor MG1655 variant (**Fig. 5b**).

We next sought to gain some insight into the observation that nine segregating variants exhibited high noise in glucose, while nine others exhibited low noise. We first tested whether the high- and low-noise phenotypes were a general phenomenon for each promoter variant, or whether the noise each variant exhibited was specific to regulation in glucose. Although we observed a strong positive correlation between noise in glucose and noise in galactose for the segregating variants (Spearman’s rho = 0.78, p = 2.4e-04), we found no correlation between noise levels in glucose and lactose (Spearman’s rho = 0.22, p = 0.39; **Fig. 6b**). This suggests that there is specific selection for high or low transcriptional noise in glucose (diversifying selection) while in lactose, directional selection has led to the filtering of mutations that increase transcriptional noise.

**Figure 6:**
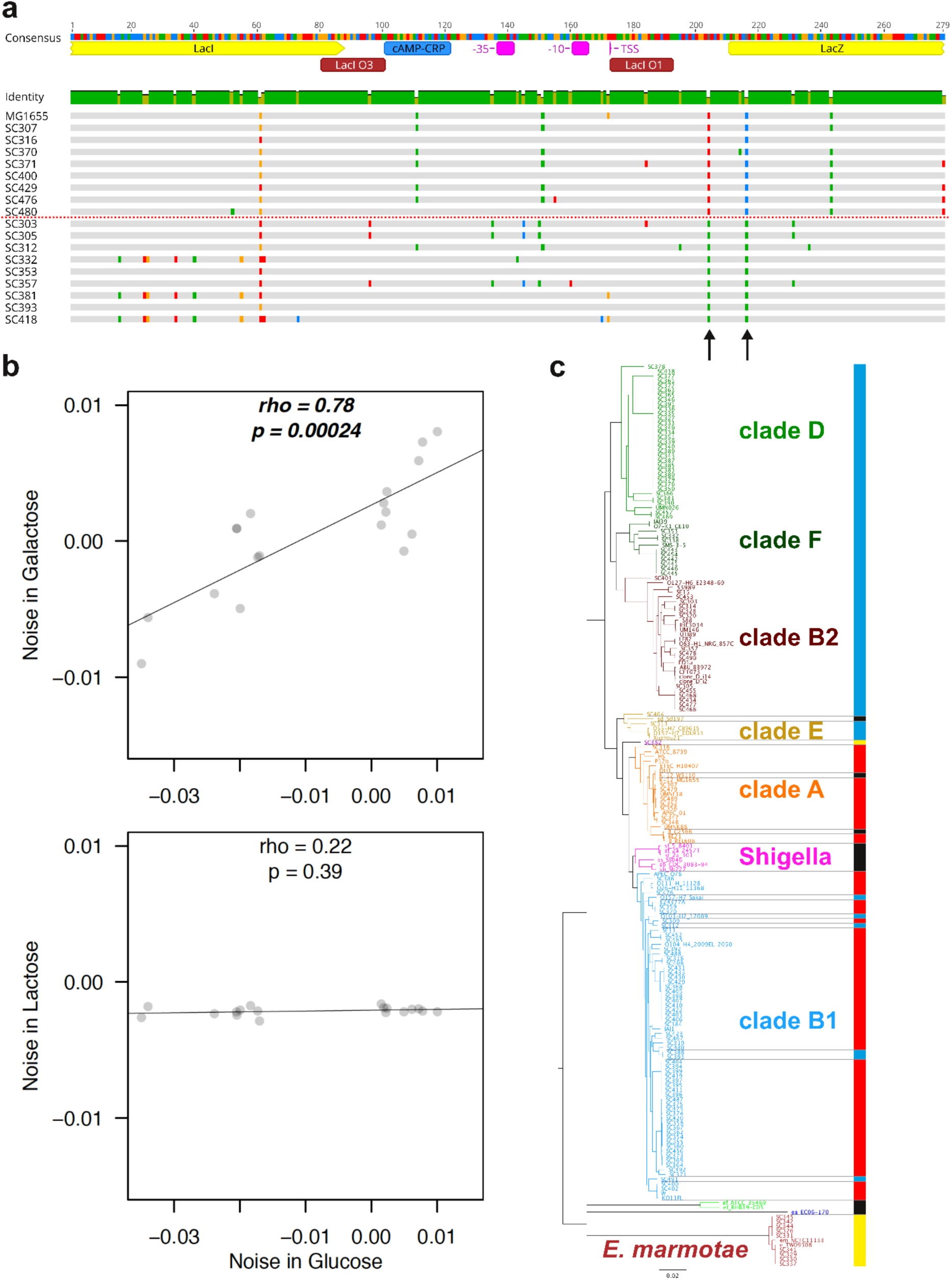
Segregating genotypes associated with high and low transcriptional noise in glucose. **a**, Alignment of segregating promoter variants. The horizontal red dotted line divides variants with high noise in glucose (above the line) and variants with low noise in glucose (below the line). The two arrows highlight the position 7 bp upstream and 6 bp downstream of the *lacZ* gene start codon. Polymorphisms in these positions are associated with high (A and C) or low noise (T at both positions). Nucleotide colors: red = A, green = T, yellow = G, blue = C, gray = matches the consensus (top sequence with annotations). **b**, Correlation between transcriptional noise in glucose and transcriptional noise in the other environments for segregating promoter variants. The black lines show linear regression of the data points. The rho and p-values were calculated using Spearman’s correlation test. **c**, Maximum likelihood phylogenetic tree constructed from the core genomes of all 135 environmental *E. coli* isolates and 56 known laboratory, type or clinical isolates of *Escherichia* species using REALPHY (Bertels et al. 2014). The colors in strain names indicate standard phylogenetic clades of the genus *Escherichia*, as indicated by the adjacent names. The bar next to the tree represents the mutational combination the strains possess at the positions highlighted by arrows in **a**: blue = TT genotype; yellow = TC genotype; red = AC genotype; black = no *lacZ* promoter found (all of these are *Shigella*, *E. fergusonii*, *E. albertii* species or laboratory *E. coli* strains with deletion of the *lac* operon). The horizontal gray lines touching the bar serve as visual aids for identifying the precise taxa.

We looked in more detail at the genotypes of the segregating *lacZ* promoter variants with high and low transcriptional noise phenotypes in glucose (**Fig. 5b**). We found that two - and only two - polymorphisms segregated perfectly between these two phenotypes. These two polymorphisms were located 7 bp upstream (-7 bp) and 6 bp downstream (+6 bp) of the *lacZ* gene start codon (**Fig. 6a**). The low noise variants had T at both positions (“TT genotypes”), while high noise variants had A and C at the respective positions (“AC genotypes”). We compared the sequences at the *lac* locus across the entire collection of 135 environmental *E. coli* isolates, and found that the frequency of TT and AC genotypes among the isolates was close to equal (62 TT vs. 63 AC). Ten of the 135 environmental *E. coli* isolates had a TC genotype. We used a core genome phylogeny to determine the distribution of these polymorphisms, and found that they segregated almost perfectly between phylogenetic groups, with all isolates in the B2, D, E, and F clades having the TT genotype associated with low noise in glucose, and the majority of isolates in the A and B1 clades having the glucose high-noise AC genotype, with six exceptions (**Fig. 6c**). These six exceptions appeared to be horizontal gene transfers from the B2, D, E, or F clades. Almost all isolates with the TC genotype were in a single highly diverged cluster of *E. marmotae*, suggesting that this genotype is ancestral, or that a rare recombination event or mutational reversion had occurred at this locus.

Notably, we found a single random variant that had changed from a high-noise AC genotype to a TC genotype, but no other random variants had changes at either the -7 bp or +6 bp positions. The -7 bp T polymorphism is associated with low-noise phenotypes, and we found that this variant exhibited a low-noise phenotype (-0.014), in contrast to the phenotype of its progenitor MG1655 variant (0.008). However, this random variant also had one additional mutation located in the LacI O3 binding site, which may also have an effect on its noise phenotype.

Nevertheless, these results provide circumstantial evidence that there are single mutations that may have strong effects on noise. Furthermore there has been long-term maintenance of genetic variants associated with either high-noise or low-noise phenotypes. This is consistent with there being balancing selection on the transcriptional noise levels conferred by *lacZ* promoter variants in a glucose environment. However, there is no strong evidence that these genotypic associations with noise phenotypes are causal, and there are a range of other reasons that could lead to the long-term maintenance of these polymorphisms, for example correlated selection on other characters, or simply chance.

### 5. Effects of genetic background on transcriptional phenotypes

Above we have quantified the *cis*-effects of segregating polymorphisms and random mutations, and have shown that in general there is directional selection acting to minimise transcriptional activity in glucose and maximise activity in lactose. We found that the vast majority of random mutations decrease plasticity relative to the MG1655 promoter variant, but there is no clear evidence that segregating promoters have been selected to maximize plasticity, perhaps due to genetic background having strong effects on plasticity. Finally, we have found evidence of diversifying selection on transcriptional noise in glucose, but directional selection in lactose.

However, these *cis*-regulatory phenotypic effects can be strongly dependent on genetic background (*trans*-regulatory effects). To gain insight into the effects of genetic background on the regulatory behavior of the *lac* operon, we transformed three of the segregating *lacZ* promoter variants into the isolates that each had originated from, i.e. their native genetic background (**Supplementary Table 1**). We also include the MG1655 promoter variant in its native MG1655 background in the analyses below.

In both glucose and galactose, we found that all variants in their native genetic background maintained the same rank order of fluorescence levels as in the non-native MG1655 background, except for a single case in galactose (**Fig. 7a**, SC418). However, the range of fluorescence levels increased considerably, with promoter variants varying more than 20-fold in fluorescence levels in glucose. This was primarily due to an almost three-fold increase in transcription for the SC312 and SC418 isolates. In strong contrast, all promoter variants converged in fluorescence levels in lactose. In the non-native genetic background, promoter variants differed by approximately three-fold; in their native genetic background, this decreased to less than two-fold, with three segregating promoter variants converging to almost exactly the same level (**Fig. 7a**). This indicates that *trans*-effects on transcriptional activity can be divergent, increasing activity in some environments (SC312 in glucose) but decreasing it in others (SC312 in lactose. These changes will necessarily affect plasticity as well.

**Figure 7:**
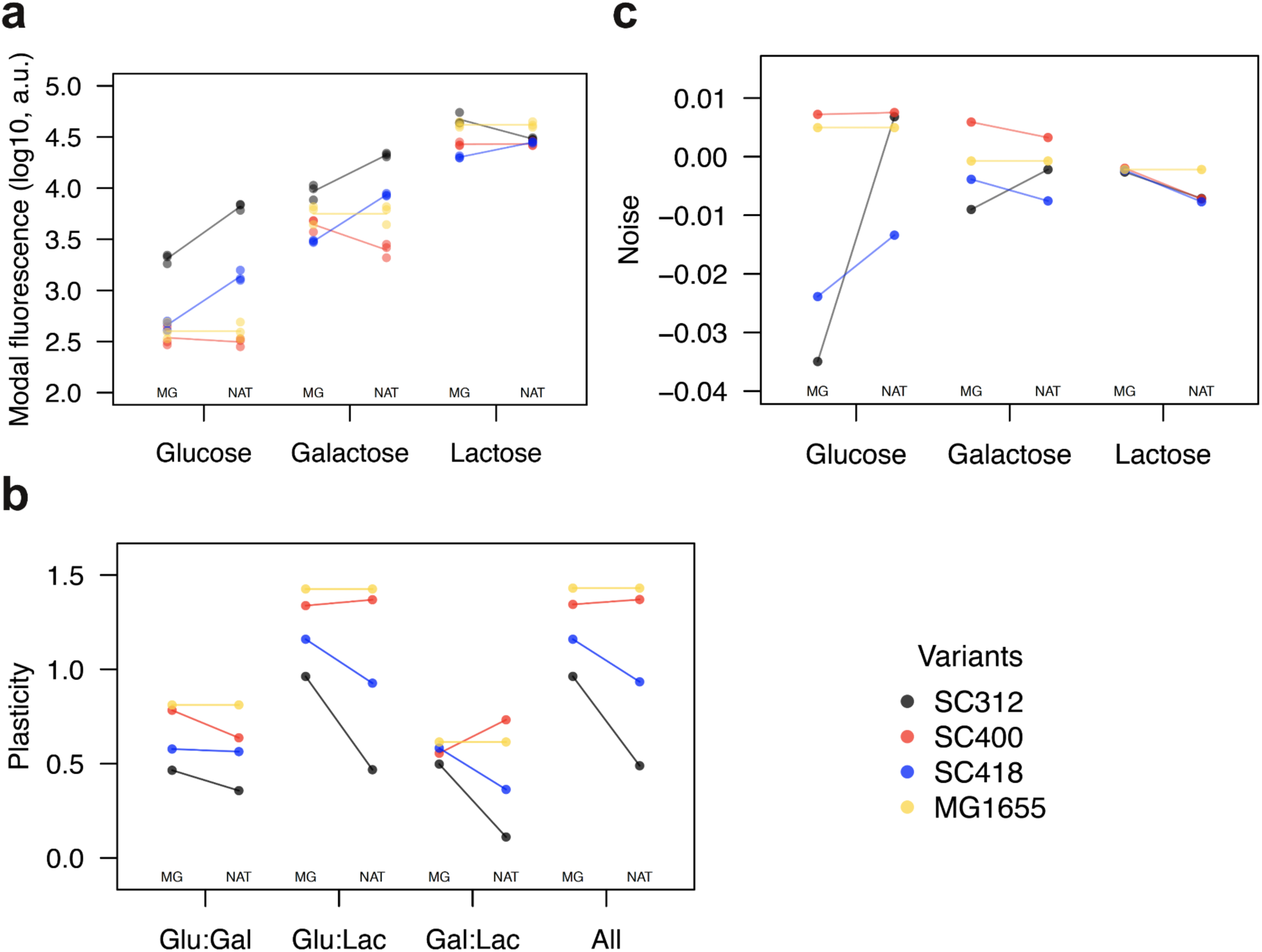
Comparison of regulatory phenotypes between segregating promoter variants in MG1655 and native genetic backgrounds. **a**, Modal population fluorescence in all three environments as indicated by x-axis labels. Each datapoint represents modal population fluorescence from one out of a total of three full biological replicates. The lines connect mean modal population fluorescence values calculated from each of the three replicates. **b**, Plasticity in all pairs of environments (Glu:Gal, Glu:Lac, and Gal:Lac) and in combination of all three environments (All). **c**, Transcriptional noise in all three environments as indicated by x-axis labels. Identical promoter variants are compared when being placed either to MG1655 genetic background (MG) or the genetic background from which each promoter variant originates (NAT). The colors distinguish promoter variants and they are connected with the same color lines within each environment or combination of. The MG1655 variant is always present in the MG1655 genetic background that serves as non-native genetic background for other variants. All the values compared between MG and NAT are thus identical.

When we considered transcriptional plasticity, we also found that in all combinations of environments the majority of promoter variants maintained the same rank order in the native genetic background as in MG1655 strain, except for SC400 in galactose and lactose (**Fig. 7b**, Gal:Lac). The maintenance of the rank order implies that the transcriptional plasticity of promoter variants follows the same pattern in both genetic backgrounds, and that from a relative standpoint, the conclusions we have derived from segregating promoter variants in the MG1655 genetic background are indicative of how the variants behave in nature. However, we also observed substantial decreases in transcriptional plasticity for SC312 and SC418 promoter variants in their native genetic backgrounds (**Fig. 7b**). This is due almost entirely to the considerable increase in transcription that occurred in glucose for these two isolates. As previously (**Fig. 4b** and **4d**) the plasticity in all three environments (**Fig. 7b**, All) primarily reflected the patterns of plasticity in glucose and lactose.

Finally, the segregating variants in their native genetic backgrounds exhibited much lower transcriptional noise in lactose as compared to random variants. This is emphasized by the fact that while there appears to be selection for low levels of noise in this environment (**Fig. 5b**, Lactose), the MG1655 variant, even when in its native background, exhibited higher levels of noise than other segregating variants. Surprisingly, in glucose, we found a very large increase in transcriptional noise of the SC312 promoter variant, which had exhibited the least noise in the MG1655 background, and which has a low-noise TT genotype (**Fig. 6a**, **6c**, and **7c**).

Interestingly, SC312 is one of six isolates in the A and B1 clades that have an apparently horizontally transferred *lac* locus (**Fig. 6c**). These results suggest that *trans*-effects (i.e. genetic background) can have strong and opposing effects to the *cis*-effects of the promoter itself.

## Discussion

Here we have investigated the actions of selection on transcriptional regulation of the canonical *lac* operon. We utilized an experimental system with a self-cleaving ribozyme RiboJ (**Fig. 1** and **Fig. 2**) that allows quantification of transcriptional effects on phenotypes while excluding translational effects (Lou et al. 2012; Vlková et al. 2021). This system also has increased sensitivity for measuring changes in transcriptional regulation (**Supplementary Figure 3**). To ensure that this system also reflects the patterns of regulatory behavior in the native *lac* locus, we replaced the native *lacZ* promoter in MG1655 with promoter variants from the natural isolates. In this experimental system, the promoter sequence can have both transcriptional and translational effects on protein expression level. Nevertheless, these results confirmed that in almost all cases, the relative transcriptional activity of each promoter in the plasmid-based system reflected the relative expression levels in the chromosome, with the rank order of fluorescence level and plasticity remaining the same in all environments except for one case, in which the promoter from isolate SC312 exhibited decreased relative fluorescence in lactose (**Supplementary Figure 4**). This may be due to low levels of LacI being bound more frequently to the single copy of the *lacZ* promoter on the chromosome, and repressing transcription, whereas on the plasmid, the LacI molecules are titrated out due to the slightly higher copy number of the promoter. However, since in the chromosomal system we also measure changes in translation it might be possible that the SC312 variant has significantly decreased translation rate as compared to the other segregating variants. We were not able to quantify noise levels for the chromosomal copies of the promoter variants, primarily because a large sample size is required to accurately calculate the noise metric. In addition, promoter variants in the chromosome exhibit fluorescence levels that are close to the limit of detection, decreasing sensitivity.

In previous work, we showed that mutations with large *cis*-effects on expression phenotypes (expression level, plasticity, and noise) are often filtered out from the *E. coli* population by both directional and stabilizing selection (Vlková and Silander 2021). However, for some regulatory phenotypes, we previously did not find evidence of selection, suggesting that changes in these phenotypes had only weak effects on fitness. While the work here is consistent with our previous results, the increased sensitivity of the assays on transcriptional phenotypes, together with additional analyses, have allowed us to discern other selective forces.

First, we have shown that there has been selection to minimize transcriptional activity in glucose and maximize activity in lactose, with random variants generally exhibiting higher transcription in glucose or lower transcription in lactose (**Fig. 3a**). Related to this, the majority of random promoter variants exhibited decreased transcriptional plasticity compared to their MG1655 progenitor (**Fig. 4**). The decreased plasticity of random variants suggests directional selection for high transcriptional plasticity in MG1655. Even so, we found no clear differences between the plasticity of segregating and random *lacZ* promoter variants. However, the segregating variants are far more diverged from MG1655 than the random variants are, still supporting the hypothesis that new mutations that decrease plasticity are filtered out by selection.

Second, using similar comparisons of segregating and random promoter variants, we have shown that selection has acted to decrease transcriptional noise in the segregating *lacZ* promoter variants in lactose. This contrasts with previous results in which we found no significant evidence of selection on noise for the *lacZ* promoter, although there was a trend for segregating variants to exhibit lower noise (Vlková and Silander 2021). The previous lack of clear evidence of selection was most likely due to a lower sample size of random variants and a lack of sensitivity in glucose.

More surprisingly, we found evidence that in glucose, segregating variants constituted two phenotypic clusters, one having high noise and the other low noise (**Fig. 5b**, Glucose). We also identified two specific polymorphisms near the start codon of the *lacZ* ORF that were associated with the two noise phenotypes (**Fig. 6a**). In addition, these two polymorphisms are ancient, having been segregating in *E. coli* since the divergence of phylogroups A and B1, in contrast to every other polymorphism segregating at the *lac* locus (**Fig. 6a** and **6c**). However, we could not establish that these two polymorphisms were causal for the noise phenotypes, nor that their long-term maintenance is due specifically to selection on noise. We did find a single random mutant that shared one of these segregating polymorphisms, and the noise phenotype of this random mutant changed considerably from the progenitor MG1655 in the direction expected if this polymorphism is causal. It has previously been shown that only a small number of genetic changes can affect noise phenotypes (Hornung et al. 2012; Metzger et al. 2015; Wolf et al. 2015; Schmiedel et al. 2019; Urchueguía et al. 2021; Vlková and Silander 2021).

Finally, we confirmed that differences in transcriptional noise measured in MG1655 strain are qualitatively similar to those observed in the native genetic backgrounds of the isolates from which the segregating variants originate, with the relative levels of noise remaining the same in most cases (**Fig. 7c**). Nevertheless, we found one clear exception, the segregating *lacZ* promoter variant from the SC312 isolate. Despite this promoter variant exhibiting the lowest noise in the MG1655 genetic background, and having the low-noise TT genotype (at positions - 7 bp and +6 bp relative to *lacZ* gene start codon) in glucose, it manifested with a high-noise phenotype when present in its original genetic background (**Fig. 7c**, SC312). Interestingly, the phylogenetic analysis suggested horizontal gene transfer (HGT) of the *lac* locus into SC312 from an isolate in the B2, D, F, or E clades. Thus, one reason for the change in the noise level of the *lac* locus in the SC312 isolate may be that there are *trans*-effects that mediate noise as well as *cis*-effects. Because of the HGT of this locus, there might be suboptimal transcriptional control of the *lac* operon in this isolate. We also observed unusually high transcriptional activity in glucose and low plasticity for the SC312 variant compared to other segregating variants (**Fig. 7a** and **7b**). We thus propose that a relatively recent HGT event has resulted in suboptimal regulation of the *lacZ* promoter. However, this suboptimal regulation may be mitigated by this isolate having evolved increased noise at the *lac* locus. Theoretical models have predicted that high noise can be beneficial when the precise expression control and plasticity is not optimal (Wolf et al. 2015; Schmiedel et al. 2019; Schmutzer and Wagner 2020). This has also been experimentally tested in *S. cerevisiae,* in which high-noise variants of the TDH3 promoter resulted in on average higher fitness as compared to low noise variants with suboptimal expression levels (Duveau et al. 2018). As we observed unusually high transcriptional activity in glucose (**Fig. 7a**) with low plasticity (**Fig. 7b**), the high transcriptional noise might be a way of mitigating the possible misregulation in glucose. Interestingly, this high noise mitigation is mediated by *trans*- rather than *cis*-genetic changes in the SC312 isolate.

## Conclusion

The data here show that for the *lacZ* promoter, natural selection acts on expression phenotypes at least in part solely through adjusting transcriptional control. We found evidence of diversifying selection acting on transcriptional noise in glucose, and two single nucleotide polymorphisms that are associated with this phenotype, but not necessarily causal. Furthermore, we found that one segregating promoter variant exhibited atypical regulatory responses both inside and outside of the genetic background of its origin. This may be a result of recent horizontal gene transfer, resulting in suboptimal regulatory phenotype. In consensus with theoretical predictions and recent experimental evidence showing that high noise can be advantageous when expression levels are suboptimal, we observed an increase in transcriptional noise in this variant. Interestingly, this increase in noise seems to be environment-specific and results from *trans*- rather than *cis*-genetic changes.

## Materials and Methods

### Construction of *lacZ* variant libraries

We created four types of *lacZ* promoter variant libraries: (1) segregating *lacZ* promoter variants placed into strain MG1655 on a plasmid; (2) random *lacZ* promoter variants placed into strain MG1655 on a plasmid; (3) segregating *lacZ* promoter variants placed into their native *E. coli* isolate on a plasmid; (4) segregating *lacZ* promoter variants placed into chromosome of strain MG1655 replacing the MG1655 promoter variant. The promoter libraries were aliquoted into 96 well microplates. Each microplate also contained a positive control consisting of the highly active murein lipoprotein (*lpp*) promoter driving GFP expression (Zaslaver et al. 2006), and a negative expression control consisting of a promoter-less plasmid pMV001 (Vlková et al. 2021). Chromosome-based library had an additional negative expression control consisting of wild-type MG1655 strain. We describe the construction of each library type separately below.

### Segregating *lacZ* variants in MG1655 genetic background

Low-copy number plasmid pMV001 with an SC101 ori, a strong RBS, self-cleaving ribozyme RiboJ and GFPmut2 gene was used for the vector backbone (Vlková et al. 2021). Both the vector backbone and segregating *lacZ* promoter variants were PCR amplified using Phusion High-Fidelity DNA polymerase with HF buffer (New England Biolabs). For promoter PCR amplification, 5 μl of pooled DNA from isolates with various variants of *lacZ* promoter was used as a DNA template (Vlková and Silander 2021). The primers for promoter amplification contained 17 nucleotide overhangs which were homologous to the ends of the vector backbone for subsequent DNA assembly. All primers used in this study are listed in **Supplementary Table 3**.

For vector PCR amplification, 0.5 ng of pMV001 plasmid DNA served as a template. After confirming a successful PCR amplification of the products on 1% agarose gel, the template DNA was digested by DpnI from the remaining reaction volume (Li et al. 2011). Both the insert and vector PCR reactions were then column-purified and we assembled the vector and promoter variants using Gibson assembly (Gibson et al. 2009) with NEBuilder® HiFi DNA Assembly Master Mix (New England Biolabs). The assembly mix was then electroporated into the electrocompetent MG1655 strain. Transformed colonies which grew on LB agar plates with 50 μg/ml Kanamycin were picked for Sanger sequencing across the insert in the vector backbone and stored as glycerol stocks. Clones with confirmed segregating promoter variants were then grown in liquid LB with Kanamycin and used to create 96 well microplate glycerol stock libraries.

### Random *lacZ* variants in MG1655 genetic background

We PCR amplified the pMV001 backbone, DpnI treated, and column-purified it the same way as described for segregating promoter variants above. We produced the promoter inserts by performing error-prone PCR using the GeneMorph II Random Mutagenesis Kit (Agilent Technologies). We used 25 ng of the plasmid construct with the MG1655 *lacZ* promoter variant cloned into it as a template DNA for the error-prone PCR to achieve approximately 1.5 SNPs per variant sequence. We used the same primers as for the segregating promoter variants (**Supplementary Table 3**). The reaction with randomly mutated promoter variants was DpnI-treated and column-purified before Gibson assembly with the NEBuilder® HiFi DNA Assembly Master Mix (New England Biolabs). The assembly mix was then electroporated into the MG1655 strain and colonies that grew on LB with Kanamycin were picked for Sanger sequencing and stored as glycerol stocks. Clones which had none or more than three SNPs in the cloned promoter insert were excluded as well as those with SNPs detected in the vector backbone (rare occasion). The rest of the clones were then re-grown in liquid LB with Kanamycin in 96 deep-well microplates overnight and 96 well microplate glycerol stock libraries were prepared from them.

### Segregating *lacZ* variants in native genetic background

We selected three environmental *E. coli* isolates: SC312, SC400, and SC418 (Ishii et al. 2006) for comparing the expression from identical segregating promoter variants inside and outside of their native genetic background (i.e., in MG1655 strain). These three isolates were selected for the different expression phenotypes produced from their *lacZ* promoter variants in the MG1655 genetic background.

We isolated the plasmids containing the three segregating *lacZ* promoter variants from the MG1655 strain using StrataPrep Plasmid Miniprep Kit (Agilent). We also isolated the promoter- less vector pMV001 the same way. We then electroporated the plasmids containing the segregating *lacZ* promoter variants into the *E. coli* isolates of the variant origin. All three isolates were also transformed with the promoter-less vector pMV001. We confirmed the presence of all desired plasmids by Sanger sequencing across the promoter insert region from clones that grew on LB with Kanamycin and aliquoted them into a microplate library.

We prepared the environmental *E. coli* isolates for electroporation as follows: we inoculated 100 ml of liquid LB in 1 l flask with 1 ml of overnight culture of environmental *E. coli* isolate (grown in liquid LB at 37°C with shaking). We then incubated the flask at 37°C with shaking (250 rpm) until the isolate reached mid-exponential phase. Once the optical density (OD600) of the media reached values between 0.5 and 0.6 we cooled down the culture by swirling the flask in ice slurry for at least 5 min and then placed it into a 4°C fridge inside the ice slurry for 1 h. Next, we spinned the cold culture at 4,200 G for 5 min at 4°C, discarded the supernatant and washed the cell pellet twice in cold 10% glycerol with centrifugation at 4,500 G for 5 min at 4°C. At the end we aliquoted 70 μl of cells in the small amount of remaining 10% glycerol into cold 1.5 ml tubes. We stored these aliquots at -80°C until use.

### Segregating *lacZ* variants in MG1655 chromosome

To modify the *lacZ* promoter of MG1655 strain at the native locus in the chromosome and obtain translational ligation of *lacZ* and GFP genes, we implemented a landing pad assay (Tas et al. 2015). In detail, we first replaced the whole *lacZ* promoter and gene, including a small part of *lacI* gene by the landing pad (TetR gene). However, using the original pTKRED helper plasmid interfered with the homologous recombination during this first step due to *lacI* gene presence in the pTKRED sequence. Because this step relies only on the λ Red system we replaced pTKRED with pKD46 helper plasmid (Datsenko and Wanner 2000). We confirmed successful landing pad integration by Sanger sequencing across the TetR insert site and heat-cured the resulting strain of the pKD46 helper plasmid using its temperature sensitive origin of replication. We constructed the first donor plasmid pMV002 so that it contained the MG1655 *lacZ* promoter variant and MG1655 *lacZ* gene translationally ligated to GFPmut2. For this purpose we first digested the synthesized insert (**Supplementary File 1**) from the plasmid on which it was delivered (Twist Biosciences) and the vector pTKDP-*neo* (Tas et al. 2015) using I-SceI restriction enzyme followed by rSAP treatment to avoid self-ligation (New England Biolabs). We then ran the digested products on 1% agarose gel and extracted the desired fragments using StrataPrep DNA Gel Extraction Kit (Agilent). We then assembled these two fragments using the NEBuilder® HiFi DNA Assembly Master Mix (New England Biolabs) with the help of four single strand oligo bridges (**Supplementary Table 3**). The other donor plasmids were constructed using the approach described above for plasmid-based libraries, just replacing the pMV001 vector with pMV002 and using a different set of primers for vector and insert PCR amplification as listed in **Supplementary Table 3**.

The donor plasmids were then transformed into the MG1655 strain having the TetR landing pad in *lac* operon and pTKRED helper plasmid. To confirm successful integration of the target sequence into the MG1655 chromosome we sequenced the whole genome of colonies that appeared blue on LB agar plates with 2 mM IPTG and 20 μg/ml X-gal after heat-curing the pTKRED helper plasmid. We then constructed a microplate library from the obtained clones.

### Flow cytometry assays

The assays and data analysis were performed the same way as previously described for the *lacZ* promoter (Vlková and Silander 2021). In short, the bacterial clones in the libraries were first inoculated into 0.5 ml of M9 minimal media with 0.4% glucose (and Kanamycin for plasmid-based libraries). After overnight growth in M9 glucose, we re-inoculated the libraries into 0.5 ml of one of the three assay media: 0.4% glucose, 0.4% galactose or 0.4% lactose. After this second overnight growth, we inoculated the libraries into the same fresh assay media into three separate microplates to obtain triplicates for each clone. The library containing segregating *lacZ* variants in the MG1655 chromosome had a different layout. This means that each triplicate was inoculated into separate wells since the first overnight in M9 with 0.4% glucose. Once the cells reached exponential growth, we diluted them into 1x PBS with ∼2.5% formaldehyde.

We performed the flow cytometry on a BD FACSCanto II machine using BD FACSDiva software version 6.1.3. We used a 488 nm laser and a 513/17 nm bandpass filter to obtain the GFP fluorescence data. We set the number of events to record from each well to 20,000. We exported the acquired data from FACSDiva software into Flow Cytometry Standard files, and performed all cell gating and fluorescence analysis using custom R scripts (flowCore package version 2.0.1; see **Supplementary Information**). We gated cells based on their maximal kernel density of forward and side scatter values, using the ellipsoidGate function from the flowCore package, and keeping about ⅓ of all events.

The modal population fluorescence was calculated as the mean of the three maximal kernel density values from the GFP fluorescence signal of three replicates. The modal coefficient of variation (mCV) was calculated separately for each of the three biological replicates (standard deviation divided by the modal population fluorescence), and the mean of these values was used as the mCV of the promoter. Replicates with fewer than 2,500 and 5,000 recorded events were excluded from the calculation of modal population fluorescence and mCV, respectively. We set the limit to 5,000 events for mCV to avoid outlier events such as machine noise to affect the mCV calculations (Vlková and Silander 2021).

When comparing *lacZ* variants that came from two separate microplate layouts, we obtained an offset for both modal population fluorescence and mCV to minimize plate-effects. We calculated these offsets as the mean of the differences between the two or three controls present in each microplate, i.e., the MG1655 promoter variant (in plasmid-based libraries only), plpp::GFPmut2 (positive control), and negative control (pMV001 or MG1655 wild-type strain). All figures contain modal fluorescence and mCV values that are corrected using these offsets. All scripts using the workflow described here, including the original data files can be found in the **Supplementary Information**.

### Quantifying transcriptional activity

To test for differences in the variation in transcriptional activity between the segregating and random variants, we calculated the modal population fluorescence values for each promoter variant in all libraries and environments as a proxy for transcriptional activity. We then tested for significant differences in variation in modal population fluorescence levels between groups using the Fligner-Killeen test of homogeneity of variances. We further tested whether increases in modal fluorescence in random variants are equally probable as decreases using two-sided binomial tests. These tests were performed against modal population fluorescence values from the MG1655 variant in each environment. The code for these calculations can be found in the **Supplementary Information**.

### Quantifying phenotypic plasticity

We calculated the phenotypic plasticity of promoter variants across all three environments as described earlier (Vlková and Silander 2021). In short, we calculated the Euclidean distance of each datapoint in three dimensions to an isocline representing null plasticity. The isocline is defined by equal values in the two or three dimensional space, i.e., x = y or x = y = z. The dimensions are defined by the modal population fluorescence values in the two or three environments compared. Each datapoint (promoter variant) is thus defined by its fluorescence values in compared environments. The closer a datapoint is to the isocline, the lower the plasticity. To test whether natural selection has acted on plasticity, we compared plasticity values from segregating and random promoter variants using a two-sided Wilcoxon rank-sum test. We also tested whether increases in plasticity in random variants are as frequent as decreases in plasticity using a two-sided binomial test. We used plasticity values from the MG1655 variant as reference for these tests in each environment combination (the code for these calculations can be found in the **Supplementary Information**).

### Quantifying transcriptional noise

We used the same metric of noise as described previously (Vlková and Silander 2021). In detail, to determine the noise within an isogenic cell population of each promoter variant, we first excluded fluorescence values from the population that were lower or higher than three standard deviations from the modal population fluorescence level. Then we calculated the mCV from the isogenic cell population as the standard deviation of the fluorescence divided by the modal population fluorescence level. We next fitted a cubic smoothing spline (smoothing parameter lambda = 0.01) to the modal population fluorescence vs. the mCV values, using all (segregating and random) promoter variants. We determined the noise levels as the difference in measured mCV from the mCV predicted from the fitted spline. We then compared the noise values between segregating and random variants using a two-sided Wilcoxon rank-sum test to determine whether selection has acted on noise. We also tested whether increase in noise due to random mutations is as likely as decrease in noise using a two-sided binomial test. Noise levels from the MG1655 variant served as a reference for the binomial test (the code for these calculations can be found in the **Supplementary Information**).

### Comparison of fluorescence values from variants with and without RiboJ

The pMV001 vector we used in this study was constructed by introducing self-cleaving ribozyme RiboJ into pUA66 vector upstream of GFPmut2 gene (Zaslaver et al. 2006; Lou et al. 2012; Vlková et al. 2021). In order to assess the changes in expression caused by RiboJ presence we used part of the dataset published earlier that contains segregating and random *lacZ* promoter variants that are identical to those used in this study, but which are cloned into pUA66 vector and thus without RiboJ (Vlková and Silander 2021).

When mapping the effect size and direction of randomly introduced SNPs we used modal population fluorescence values only from those random variants which contained a single SNP relative to the MG1655 *lacZ* promoter variant. Information about the TF binding sites, -10 and - 35 elements, and ORFs was taken from the EcoCyc database (Karp et al. 2018), and only the annotations associated with σ^70^ driven TSS were used. Using the Sanger sequencing results, we identified the location of all SNPs for each random variant. We also compared the modal population fluorescence values from all variants shared between pUA66 and pMV001-based systems using Spearman’s correlation test (the code for all calculations can be found in the **Supplementary Information**).

## Acknowledgements

We thank Tim Cooper and Andrea Sajuthi for valuable comments on the final draft of this manuscript. This work was supported by a Marsden Grant (grant MAU1703) awarded to OKS. The funder had no role in study design, data collection and interpretation, or the decision to submit the work for publication.

## Author contributions

MV and OKS conceived the project, designed the experiments and analyses and wrote the paper. OKS supervised the project. MV performed all experiments and analyses.

## Data availability

The original data files and scripts with data analyses that support the findings of this study are available in the **Supplementary Information** of this study.

## Supplementary Information

All scripts with access to original data files can access through https://doi.org/10.5281/zenodo.6364515, the Supplementary Files and all Figures in high quality can be accessed through Figshare DOI: https://doi.org/10.6084/m9.figshare.c.5624566.v1.

**Supplementary Figure 1:**
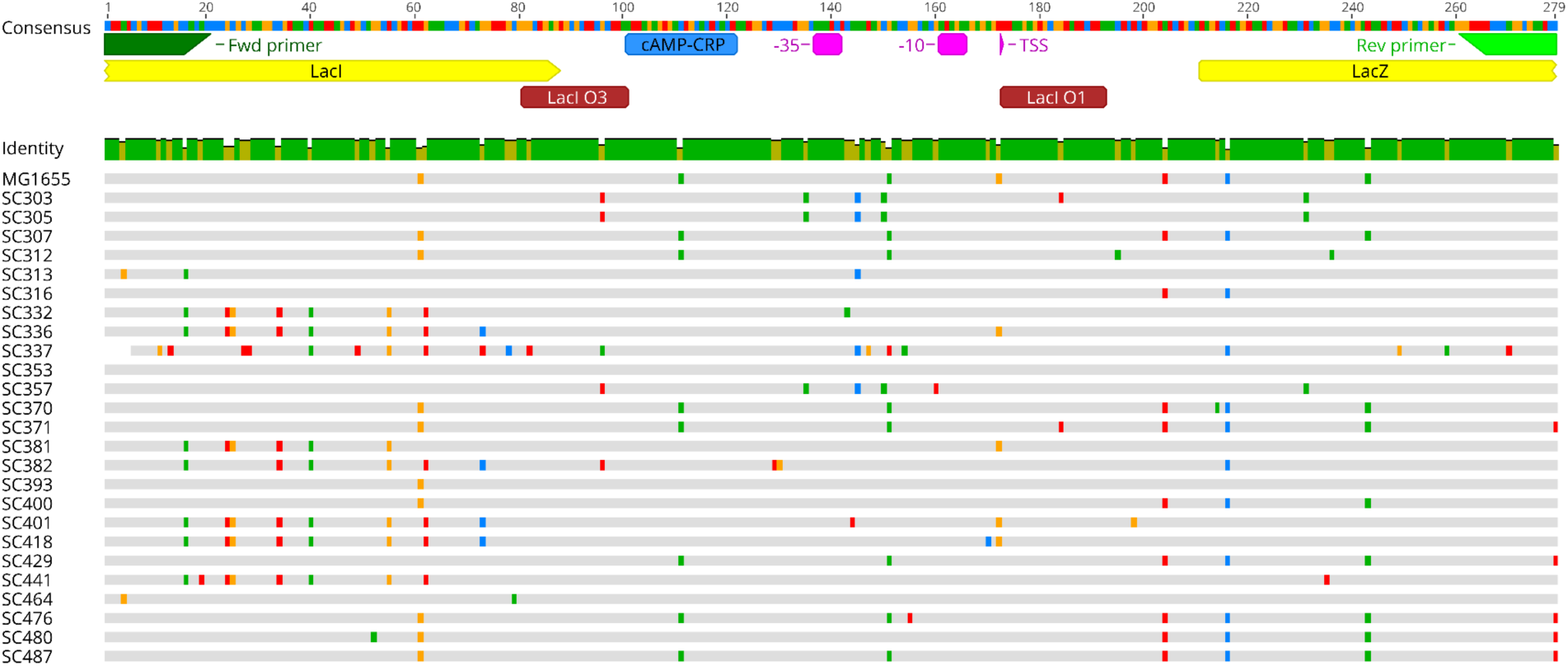
Sequence alignment of all segregating promoter variants found among 135 environmental *E. coli* isolates and MG1655 laboratory strain. Nucleotide colors: red = A, green = T, yellow = G, blue = C, gray = matches the consensus (top sequence with annotations). The green primer annotations indicate the primers used for PCR amplification and assembly of the variants with the pMV001 vector (**Fig. 1d**).

**Supplementary Table 1:** List of segregating *lacZ* promoter variants. The variant names correspond to representative isolates in which the variants were identified. The same variant naming is used throughout the manuscript. Bolded variants were cloned into both pMV001 vector and the pUA66 vector (lacking RiboJ), and used for comparison of fluorescence in the **Supplementary Figure 3**. * These variants were also cloned into the MG1655 chromosome and were assayed both in MG1655 and in the isolate of their origin. † Segregating variants that were not cloned.

| Variant | Polymorphisms relative to MG1655 variant | Present in N isolates out of 135 |
| --- | --- | --- |
| **MG1655\*** | 0 | 0 |
| **SC307** | 1 | 17 |
| **SC370** | 2 | 2 |
| **SC371** | 3 | 1 |
| **SC400\*** | 3 | 7 |
| **SC429** | 3 | 4 |
| **SC476** | 3 | 1 |
| **SC316** | 5 | 1 |
| **SC480** | 5 | 1 |
| **SC312\*** | 6 | 2 |
| SC393 | 6 | 2 |
| SC353 | 7 | 1 |
| **SC305** | 12 | 12 |
| **SC381** | 12 | 2 |
| **SC303** | 13 | 1 |
| **SC357** | 13 | 1 |
| **SC332** | 15 | 2 |
| **SC418\*** | 15 | 8 |
| SC487† | 2 | 29 |
| SC464† | 9 | 1 |
| SC313† | 10 | 1 |
| SC336† | 14 | 22 |
| SC382† | 15 | 1 |
| SC401† | 16 | 1 |
| SC441† | 16 | 6 |
| SC337† | 19 | 9 |

**Supplementary Figure 2:**
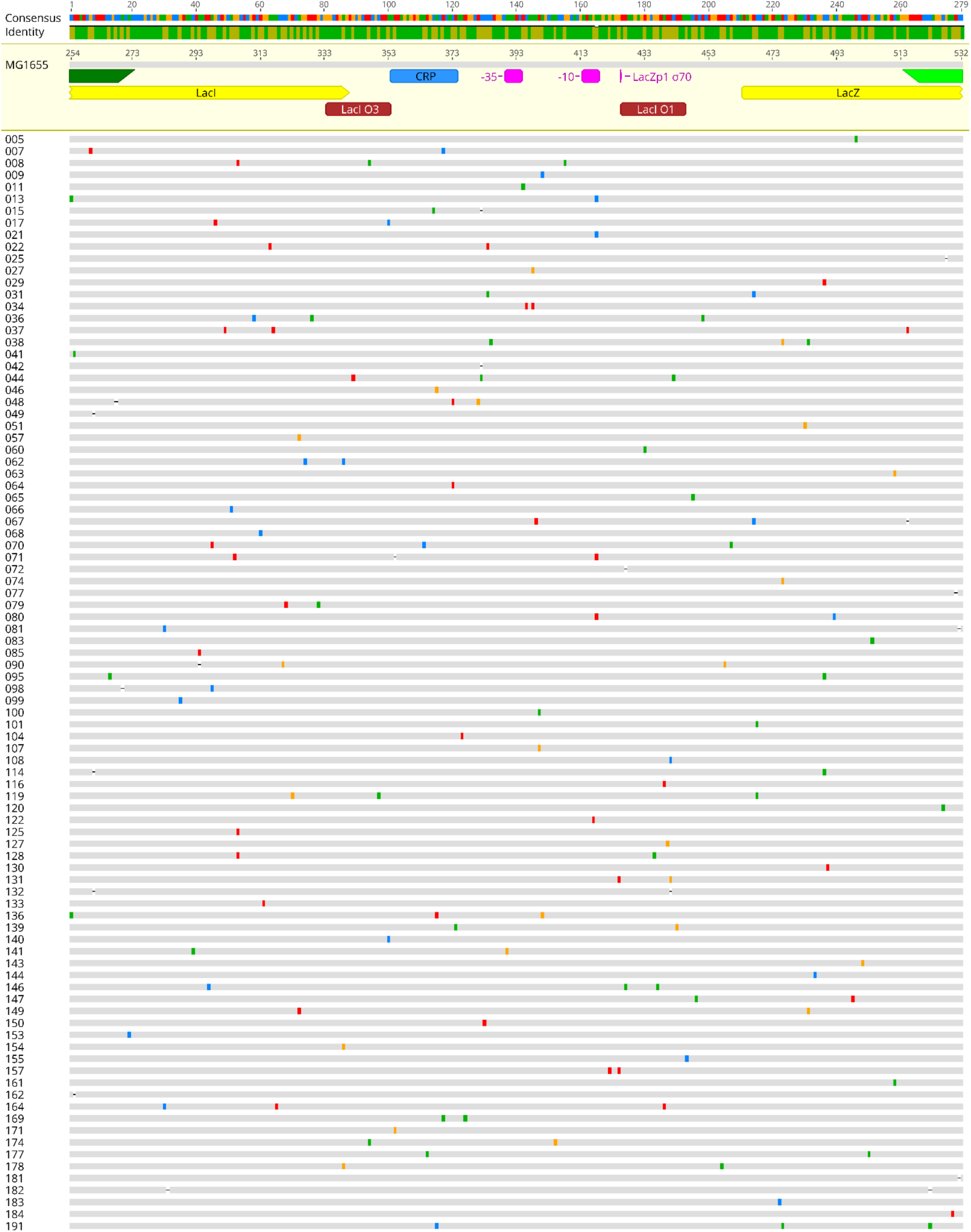
Sequence alignment of random promoter variants. Nucleotide colors: red = A, green = T, yellow = G, blue = C, gray = same as the MG1655 reference variant from which the random variants are derived (the sequence with annotations). The dashes in the sequences indicate indels.

**Supplementary Table 2:** List of random *lacZ* promoter variants. The same variant naming is used throughout the manuscript. The variants in bold were also cloned into the pUA66 vector and used for comparison of fluorescence in the **Supplementary Figure 3**.

| Variant | Mutations relative to the MG1655 variant |
| --- | --- |
| **005** | 1 |
| 009 | 1 |
| 011 | 1 |
| **021** | 1 |
| 025 | 1 |
| 027 | 1 |
| **029** | 1 |
| 041 | 1 |
| 042 | 1 |
| 046 | 1 |
| 049 | 1 |
| 051 | 1 |
| 057 | 1 |
| 060 | 1 |
| **063** | 1 |
| 064 | 1 |
| **065** | 1 |
| 066 | 1 |
| 068 | 1 |
| 072 | 1 |
| **074** | 1 |
| 077 | 1 |
| 083 | 1 |
| 085 | 1 |
| 099 | 1 |
| **100** | 1 |
| 101 | 1 |
| **104** | 1 |
| **107** | 1 |
| 108 | 1 |
| **116** | 1 |
| 120 | 1 |
| **122** | 1 |
| 125 | 1 |
| **127** | 1 |
| 130 | 1 |
| **133** | 1 |
| 140 | 1 |
| **143** | 1 |
| 144 | 1 |
| 150 | 1 |
| 153 | 1 |
| **154** | 1 |
| 155 | 1 |
| 161 | 1 |
| 162 | 1 |
| **171** | 1 |
| 181 | 1 |
| 183 | 1 |
| 184 | 1 |
| 007 | 2 |
| 013 | 2 |
| 015 | 2 |
| 017 | 2 |
| 022 | 2 |
| 031 | 2 |
| 034 | 2 |
| **062** | 2 |
| 079 | 2 |
| 080 | 2 |
| 081 | 2 |
| 095 | 2 |
| 098 | 2 |
| 114 | 2 |
| 128 | 2 |
| **131** | 2 |
| 132 | 2 |
| 139 | 2 |
| 141 | 2 |
| 147 | 2 |
| **149** | 2 |
| 157 | 2 |
| 169 | 2 |
| **174** | 2 |
| **177** | 2 |
| 178 | 2 |
| 182 | 2 |
| 008 | 3 |
| **036** | 3 |
| 037 | 3 |
| 038 | 3 |
| 044 | 3 |
| 048 | 3 |
| 067 | 3 |
| 070 | 3 |
| 071 | 3 |
| 090 | 3 |
| 119 | 3 |
| 136 | 3 |
| **146** | 3 |
| 164 | 3 |
| 191 | 3 |

**Supplementary Figure 3:**
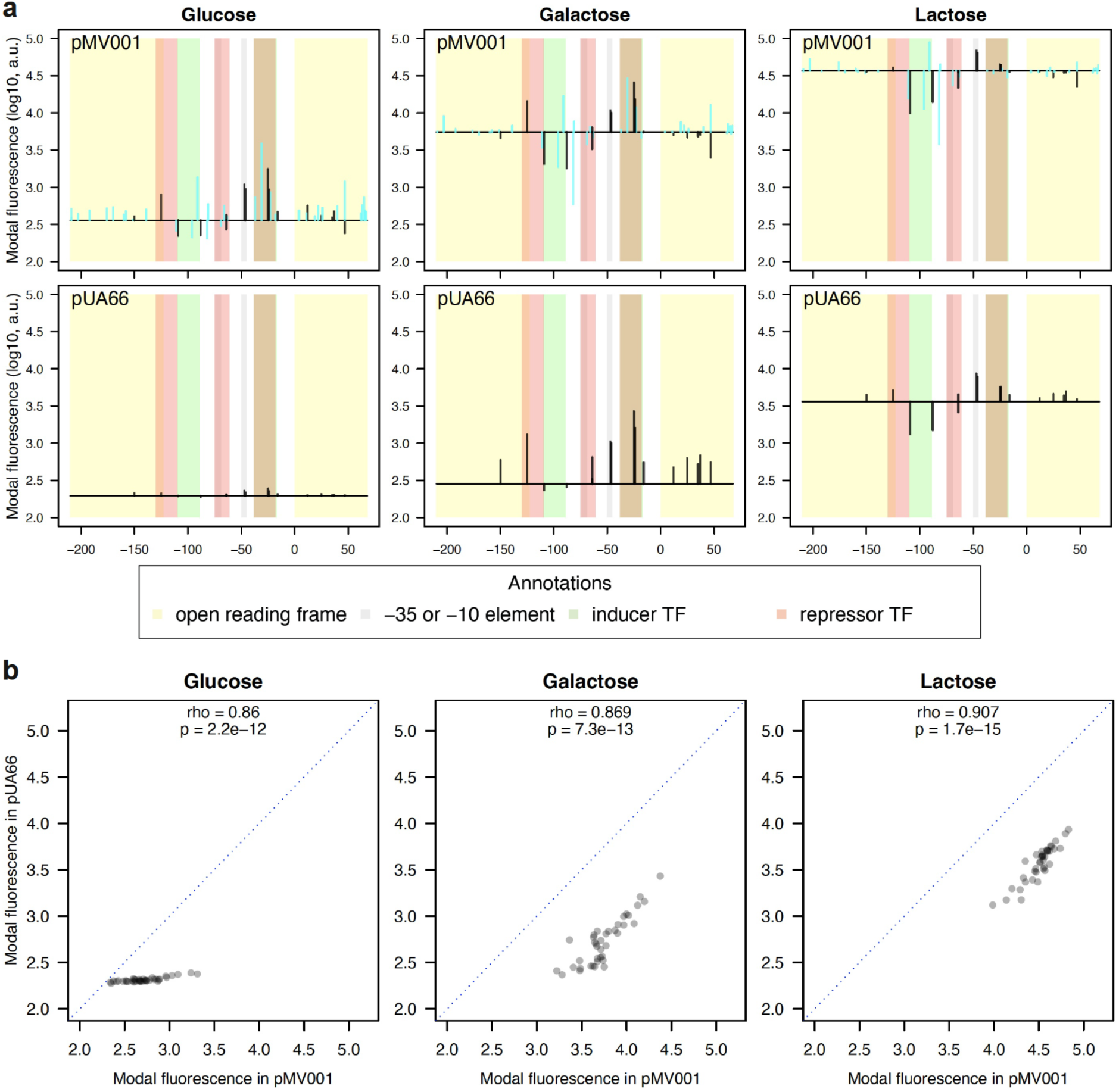
The changes in fluorescence due to random mutations are correlated between the two plasmid systems. **a**, Changes in fluorescence from the MG1655 promoter variant due to single random mutations for all environments and both plasmid systems (pUA66 and pMV001). The x-axes indicate the position relative to the *lacZ* gene start codon. The horizontal black line indicates the fluorescence level of the MG1655 *lacZ* promoter variant in that environment and with that plasmid system. The vertical lines show the direction and size of the change in fluorescence level when the MG1655 promoter variant is mutated at that specific position. The black vertical lines show mutation changes that are present in both plasmid systems, while the cyan vertical lines are mutations specific just for the dataset with RiboJ. The brown stripes result from overlapping repressor and activator TF binding sites. **b**, Comparison of modal fluorescence from all variants which are shared by both plasmid systems in MG1655 genetic background. The modal fluorescence levels are strongly correlated in all environments. Each data point in **a** and **b** is a result of three full biological replicates (**Materials and Methods**). The blue dotted lines indicate identical modal fluorescence in both plasmid systems (x = y). The rho and p-values were calculated using Pearson’s correlation test.

**Supplementary Table 3:**
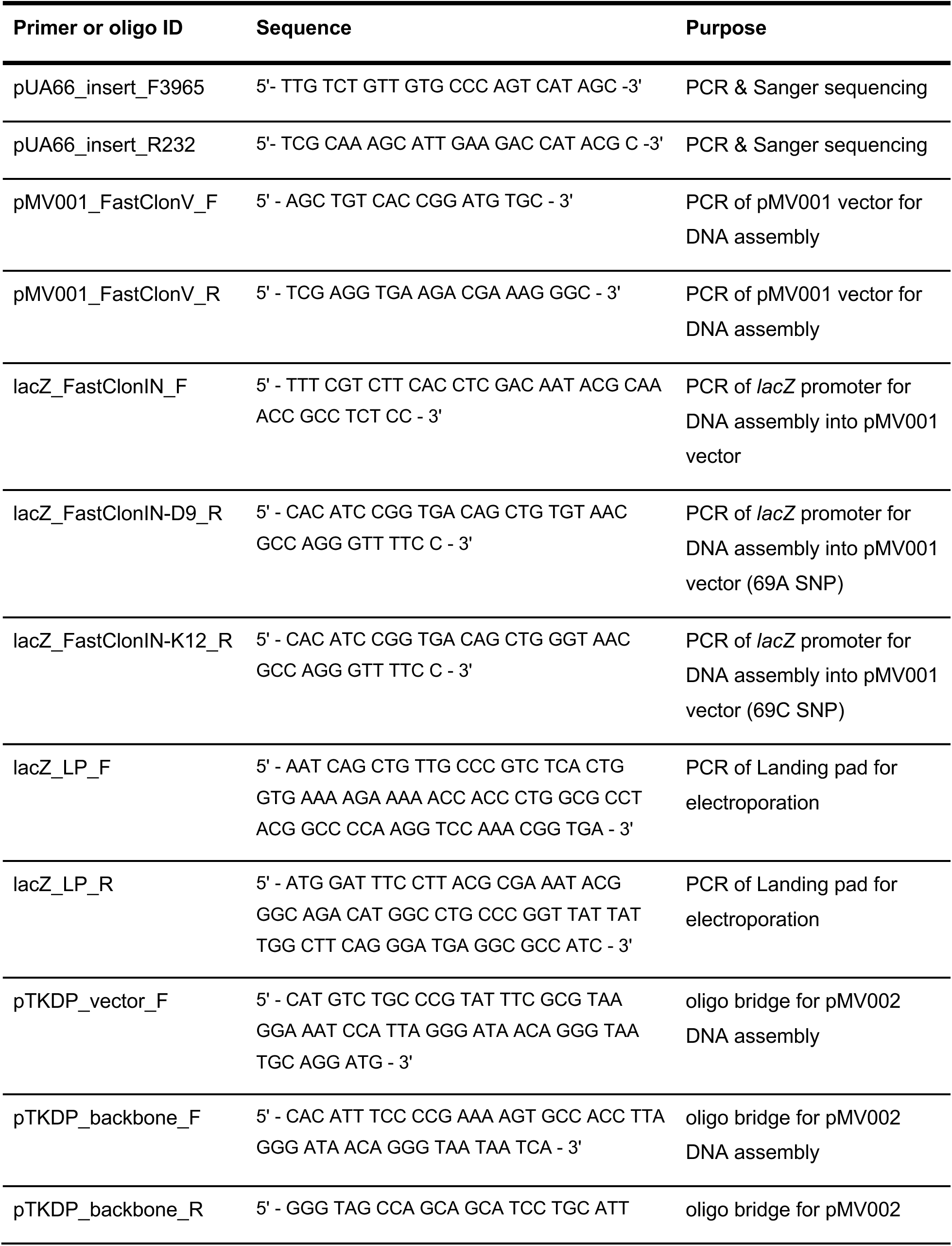
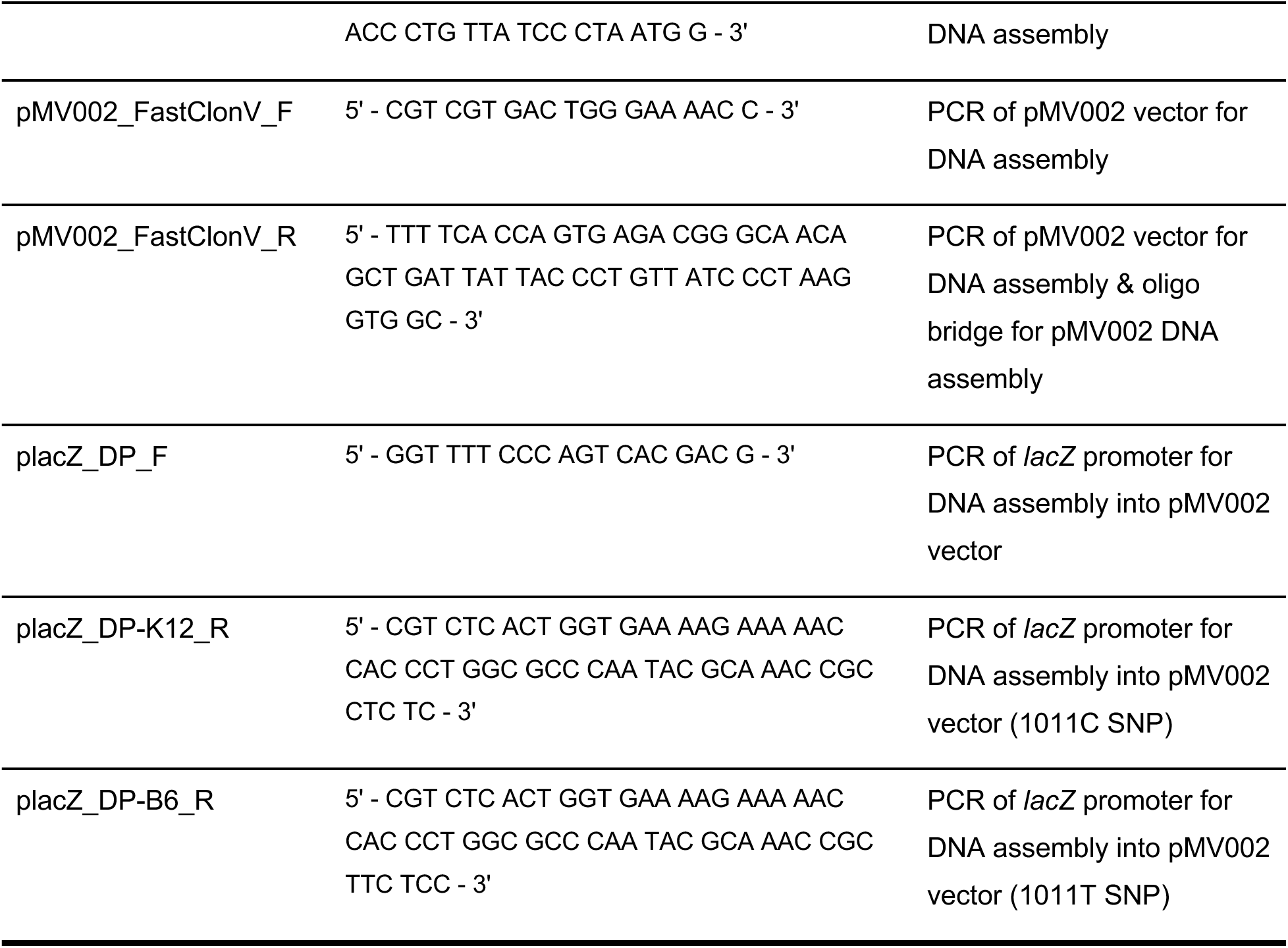
Primers and oligos used in this work. There are two versions of the reverse primer sets due to a SNP presence in the priming area of the *lacZ* gene (C69A) and *lacI* gene (C1011T).

**Supplementary Figure 4:**
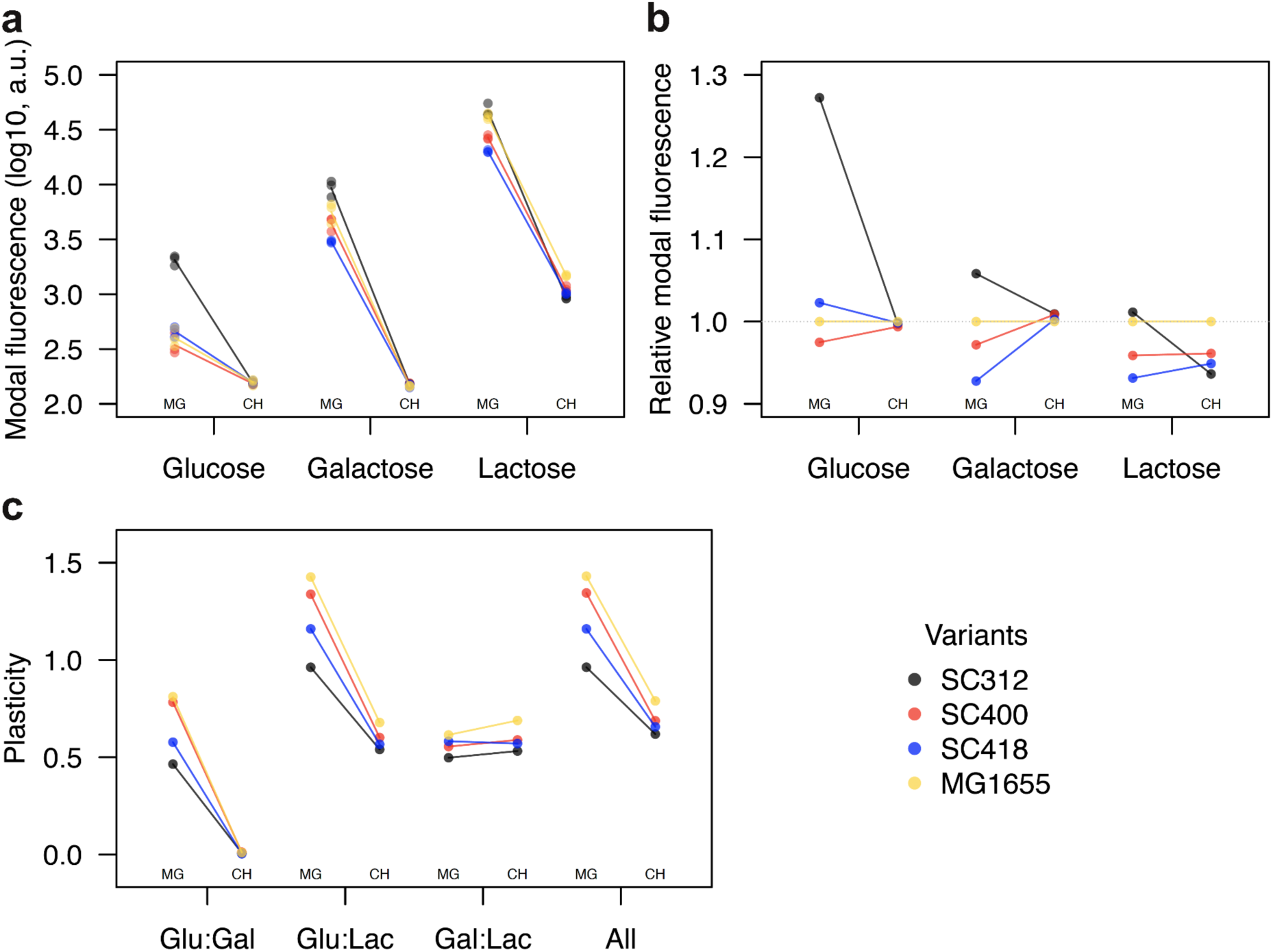
Comparison of regulatory phenotypes between segregating promoter variants in MG1655 on a plasmid and in the chromosome. **a**, Modal population fluorescence in all three environments as indicated by x-axis labels. Each datapoint represents modal population fluorescence from one out of a total of three full biological replicates. The lines connect mean modal population fluorescence values calculated from each of the three replicates. **b**, Modal population fluorescence relative to the MG1655 variant in all three environments as indicated by x-axis labels. **c**, Plasticity in all pairs of environments (Glu:Gal, Glu:Lac, and Gal:Lac) and in combination of all three environments (All). Identical promoter variants are compared when being placed either to MG1655 genetic background on a plasmid (MG) or into the *lac* operon on chromosome (CH). The colors distinguish promoter variants and they are connected with the same color lines within each environment or combination of.

## References

1. Bailey SF, Alonso Morales LA, Kassen R. 2021. Effects of synonymous mutations beyond codon bias: The evidence for adaptive 4 synonymous substitutions from microbial evolution experiments. Genome Biol Evol. 13(9):evab141. doi:10.1093/gbe/evab141.

2. Bertels F, Silander OK, Pachkov M, Rainey PB, van Nimwegen E. 2014. Automated Reconstruction of Whole-Genome Phylogenies from Short-Sequence Reads. Mol Biol Evol. 31(5):1077–1088. doi:10.1093/molbev/msu088.

3. Browning DF, Busby SJW. 2016. Local and global regulation of transcription initiation in bacteria. Nat Rev Microbiol. 14(10):638–650. doi:10.1038/nrmicro.2016.103.

4. Carrier TA, Keasling JD. 1997. Engineering mRNA Stability in E. coli by the Addition of Synthetic Hairpins Using a 5’ Cassette System. Biotechnol Bioeng. 55(3):577–580. doi:10.1002/(SICI)1097-0290(19970805)55:3<577::AID-BIT16>3.0.CO;2-D.

5. Clifton KP, Jones EM, Paudel S, Marken JP, Monette CE, Halleran AD, Epp L, Saha MS. 2018. The genetic insulator RiboJ increases expression of insulated genes. J Biol Eng. 12(1):23. doi:10.1186/s13036-018-0115-6.

6. Datsenko KA, Wanner BL. 2000. One-step inactivation of chromosomal genes in Escherichia coli K-12 using PCR products. Proc Natl Acad Sci. 97(12):6640–6645. doi:10.1073/pnas.120163297.

7. Desnoyers G, Bouchard M-P, Massé E. 2013. New insights into small RNA-dependent translational regulation in prokaryotes. Trends Genet. 29(2):92–98. doi:10.1016/j.tig.2012.10.004.

8. Duveau F, Hodgins-Davis A, Metzger BP, Yang B, Tryban S, Walker EA, Lybrook T, Wittkopp PJ. 2018. Fitness effects of altering gene expression noise in Saccharomyces cerevisiae. eLife. 7:e37272. doi:10.7554/eLife.37272.

9. Duveau F, Yuan DC, Metzger BPH, Hodgins-Davis A, Wittkopp PJ. 2017. Effects of mutation and selection on plasticity of a promoter activity in Saccharomyces cerevisiae. Proc Natl Acad Sci. 114(52):E11218–E11227. doi:10.1073/pnas.1713960115.

10. Frumkin I, Schirman D, Rotman A, Li F, Zahavi L, Mordret E, Asraf O, Wu S, Levy SF, Pilpel Y. 2017. Gene Architectures that Minimize Cost of Gene Expression. Mol Cell. 65(1):142–153. doi:10.1016/j.molcel.2016.11.007.

11. Gibson DG, Young L, Chuang R-Y, Venter JC, Hutchison CA, Smith HO. 2009. Enzymatic assembly of DNA molecules up to several hundred kilobases. Nat Methods. 6(5):343–345. doi:10.1038/nmeth.1318.

12. Gresham D, Desai MM, Tucker CM, Jenq HT, Pai DA, Ward A, DeSevo CG, Botstein D, Dunham MJ. 2008. The Repertoire and Dynamics of Evolutionary Adaptations to Controlled Nutrient-Limited Environments in Yeast. Snyder M, editor. PLoS Genet. 4(12):e1000303. doi:10.1371/journal.pgen.1000303.

13. Hawkins JS, Silvis MR, Koo B-M, Peters JM, Osadnik H, Jost M, Hearne CC, Weissman JS, Todor H, Gross CA. 2020. Mismatch-CRISPRi Reveals the Co-varying Expression-Fitness Relationships of Essential Genes in Escherichia coli and Bacillus subtilis. Cell Syst. 11(5):523–535. doi:10.1016/j.cels.2020.09.009.

14. Hodgins-Davis A, Duveau F, Walker EA, Wittkopp PJ. 2019. Empirical measures of mutational effects define neutral models of regulatory evolution in Saccharomyces cerevisiae. Proc Natl Acad Sci. 116(42):21085–21093. doi:10.1073/pnas.1902823116.

15. Hornung G, Bar-Ziv R, Rosin D, Tokuriki N, Tawfik DS, Oren M, Barkai N. 2012. Noise-mean relationship in mutated promoters. Genome Res. 22(12):2409–2417. doi:10.1101/gr.139378.112.

16. Hudson JM, Fried MG. 1990. Co-operative Interactions Between the Catabolite Gene Activator Protein and the lac Repressor at the Lactose Promoter. J Mol Biol. 214(2):381–396. doi:10.1016/0022-2836(90)90188-R.

17. Ishii S, Ksoll WB, Hicks RE, Sadowsky MJ. 2006. Presence and Growth of Naturalized Escherichia coli in Temperate Soils from Lake Superior Watersheds. Appl Environ Microbiol. 72(1):612–621. doi:10.1128/AEM.72.1.612-621.2006.

18. Iyer V, Struhl K. 1996. Absolute mRNA levels and transcriptional initiation rates in Saccharomyces cerevisiae. Proc Natl Acad Sci. 93(11):5208–5212. doi:10.1073/pnas.93.11.5208.

19. Jobe A, Bourgeois S. 1972. lac Repressor-Operator Interaction: VI. The natural inducer of the lac operon. J Mol Biol. 69(3):397–404. doi:10.1016/0022-2836(72)90253-7.

20. Karp PD, Ong WK, Paley S, Billington R, Caspi R, Fulcher C, Kothari A, Krummenacker M, Latendresse M, Midford PE, et al. 2018. The EcoCyc Database. EcoSal Plus. 6(1). doi:10.1128/ecosalplus.ESP-0006-2018.

21. Kennell D, Riezman H. 1977. Transcription and translation initiation frequencies of the Escherichia coli lac operon. J Mol Biol. 114(1):1–21. doi:10.1016/0022-2836(77)90279-0.

22. Keren L, Hausser J, Lotan-Pompan M, Vainberg Slutskin I, Alisar H, Kaminski S, Weinberger A, Alon U, Milo R, Segal E. 2016. Massively Parallel Interrogation of the Effects of Gene Expression Levels on Fitness. Cell. 166(5):1282–1294.e18. doi:10.1016/j.cell.2016.07.024.

23. Khademi SMH, Sazinas P, Jelsbak L. 2019. Within-Host Adaptation Mediated by Intergenic Evolution in Pseudomonas aeruginosa. Golding B, editor. Genome Biol Evol. 11(5):1385– 1397. doi:10.1093/gbe/evz083.

24. Kudla G, Murray AW, Tollervey D, Plotkin JB. 2009. Coding-Sequence Determinants of Gene Expression in Escherichia coli. Science. 324(5924):255–258. doi:10.1126/science.1170160.

25. Kuhlman T, Zhang Z, Saier MH, Hwa T. 2007. Combinatorial transcriptional control of the lactose operon of Escherichia coli. Proc Natl Acad Sci. 104(14):6043–6048. doi:10.1073/pnas.0606717104.

26. Kuo J-T, Chang Y-J, Tseng C-P. 2003. Growth rate regulation of lac operon expression in Escherichia coli is cyclic AMP dependent. FEBS Lett. 553(3):397–402. doi:10.1016/S0014-5793(03)01071-8.

27. Li C, Wen A, Shen B, Lu J, Huang Y, Chang Y. 2011. FastCloning: a highly simplified, purification-free, sequence- and ligation-independent PCR cloning method. BMC Biotechnol. 11(1):92. doi:10.1186/1472-6750-11-92.

28. Lou C, Stanton B, Chen Y-J, Munsky B, Voigt CA. 2012. Ribozyme-based insulator parts buffer synthetic circuits from genetic context. Nat Biotechnol. 30(11):1137–1142. doi:10.1038/nbt.2401.

29. Maeda YT, Sano M. 2006. Regulatory Dynamics of Synthetic Gene Networks with Positive Feedback. J Mol Biol. 359(4):1107–1124. doi:10.1016/j.jmb.2006.03.064.

30. Metzger BPH, Yuan DC, Gruber JD, Duveau F, Wittkopp PJ. 2015. Selection on noise constrains variation in a eukaryotic promoter. Nature. 521(7552):344–347. doi:10.1038/nature14244.

31. Mustaev A, Roberts J, Gottesman M. 2017. Transcription elongation. Transcription. 8(3):150– 161. doi:10.1080/21541264.2017.1289294.

32. Naville M, Gautheret D. 2009. Transcription attenuation in bacteria: theme and variations. Brief Funct Genomic Proteomic. 8(6):482–492. doi:10.1093/bfgp/elp025.

33. Neves D, Vos S, Blank LM, Ebert BE. 2020. Pseudomonas mRNA 2.0: Boosting Gene Expression Through Enhanced mRNA Stability and Translational Efficiency. Front Bioeng Biotechnol. 7:458. doi:10.3389/fbioe.2019.00458.

34. Oehler S, Eismann E, Krämer H, Müller-Hill B. 1990. The three operators of the lac operon cooperate in repression. EMBO J. 9(4):973–979. doi:10.1002/j.1460-2075.1990.tb08199.x.

35. Ozbudak EM, Thattai M, Lim HN, Shraiman BI, van Oudenaarden A. 2004. Multistability in the lactose utilization network of Escherichia coli. Nature. 427(6976):737–740. doi:10.1038/nature02298.

36. Phillips KN, Widmann S, Lai H-Y, Nguyen J, Ray JCJ, Balázsi G, Cooper TF. 2019. Diversity in lac Operon Regulation among Diverse Escherichia coli Isolates Depends on the Broader Genetic Background but Is Not Explained by Genetic Relatedness. Cooper VS, Whiteley M, editors. mBio. 10(6):e02232–19. doi:10.1128/mBio.02232-19.

37. Rech GE, Sanz-Martín JM, Anisimova M, Sukno SA, Thon MR. 2014. Natural Selection on Coding and Noncoding DNA Sequences Is Associated with Virulence Genes in a Plant Pathogenic Fungus. Genome Biol Evol. 6(9):2368–2379. doi:10.1093/gbe/evu192.

38. Sayut DJ, Kambam PKR, Sun L. 2007. Noise and kinetics of LuxR positive feedback loops. Biochem Biophys Res Commun. 363(3):667–673. doi:10.1016/j.bbrc.2007.09.057.

39. Schmiedel JM, Carey LB, Lehner B. 2019. Empirical mean-noise fitness landscapes reveal the fitness impact of gene expression noise. Nat Commun. 10(1):3180. doi:10.1038/s41467-019-11116-w.

40. Schmutzer M, Wagner A. 2020. Gene expression noise can promote the fixation of beneficial mutations in fluctuating environments. Goulian M, editor. PLOS Comput Biol. 16(10):e1007727. doi:10.1371/journal.pcbi.1007727.

41. Silander OK, Nikolic N, Zaslaver A, Bren A, Kikoin I, Alon U, Ackermann M. 2012. A Genome-Wide Analysis of Promoter-Mediated Phenotypic Noise in Escherichia coli. Matic I, editor. PLoS Genet. 8(1):e1002443. doi:10.1371/journal.pgen.1002443.

42. Taniguchi Y, Choi PJ, Li G-W, Chen H, Babu M, Hearn J, Emili A, Xie XS. 2010. Quantifying E. coli Proteome and Transcriptome with Single-Molecule Sensitivity in Single Cells. Science. 329(5991):533–538. doi:10.1126/science.1188308.

43. Tas H, Nguyen CT, Patel R, Kim NH, Kuhlman TE. 2015. An Integrated System for Precise Genome Modification in Escherichia coli. Herman C, editor. PLOS ONE. 10(9):e0136963. doi:10.1371/journal.pone.0136963.

44. Tenaillon O, Barrick JE, Ribeck N, Deatherage DE, Blanchard JL, Dasgupta A, Wu GC, Wielgoss S, Cruveiller S, Médigue C, et al. 2016. Tempo and mode of genome evolution in a 50,000-generation experiment. Nature. 536(7615):165–170. doi:10.1038/nature18959.

45. Urchueguía A, Galbusera L, Chauvin D, Bellement G, Julou T, van Nimwegen E. 2021. Genome-wide gene expression noise in Escherichia coli is condition-dependent and determined by propagation of noise through the regulatory network. Balaban N, editor. PLOS Biol. 19(12):e3001491. doi:10.1371/journal.pbio.3001491.

46. Vlková M, Morampalli BR, Silander OK. 2021. Efficiency of the synthetic self-splicing RiboJ ribozyme is robust to cis- and trans-changes in genetic background. MicrobiologyOpen. 10(4):e1232. doi:10.1002/mbo3.1232.

47. Vlková M, Silander OK. 2021. Gene regulation is commonly selected for high plasticity and low noise. bioRxiv. doi:10.1101/2021.07.18.452581.

48. Wanner BL, Kodaira R, Neidhardt FC. 1978. Regulation of lac Operon Expression: Reappraisal of the Theory of Catabolite Repression. J Bacteriol. 136(3):947–954. doi:10.1128/jb.136.3.947-954.1978.

49. Wheatley RW, Lo S, Jancewicz LJ, Dugdale ML, Huber RE. 2013. Structural Explanation for Allolactose (lac Operon Inducer) Synthesis by lacZ β-Galactosidase and the Evolutionary Relationship between Allolactose Synthesis and the lac Repressor. J Biol Chem. 288(18):12993–13005. doi:10.1074/jbc.M113.455436.

50. Whitaker WR, Lee H, Arkin AP, Dueber JE. 2015. Avoidance of Truncated Proteins from Unintended Ribosome Binding Sites within Heterologous Protein Coding Sequences. ACS Synth Biol. 4(3):249–257. doi:10.1021/sb500003x.

51. Wolf L, Silander OK, van Nimwegen E. 2015. Expression noise facilitates the evolution of gene regulation. eLife. 4:e05856. doi:10.7554/eLife.05856.

52. Yona AH, Alm EJ, Gore J. 2018. Random sequences rapidly evolve into de novo promoters. Nat Commun. 9(1):1530. doi:10.1038/s41467-018-04026-w.

53. Zaslaver A, Bren A, Ronen M, Itzkovitz S, Kikoin I, Shavit S, Liebermeister W, Surette MG, Alon U. 2006. A comprehensive library of fluorescent transcriptional reporters for Escherichia coli. Nat Methods. 3(8):623–628. doi:10.1038/nmeth895.

